# Multi-model Segmentation and Morphometric Quantification of Cerebral Amyloid Angiopathy in Alzheimer’s Disease Whole Slide Histopathology Images

**DOI:** 10.64898/2026.07.16.739032

**Authors:** Hossein Tahmasebidehkordi, Afshin Bahramy, Dana R. Julian, Jonathan A. Cohen, Makayla Neal, V. K. Cody Bumgardner, Peter T. Nelson, Thomas M. Pearce, Julia Kofler

## Abstract

**Introduction:** Cerebral amyloid angiopathy (CAA) is characterized by amyloid-beta deposition in cortical and leptomeningeal vessels and associated with cognitive impairment and hemorrhage. Current neuropathological assessments rely on semiquantitative grading and lack vessel-level resolution and scalability. Existing computational pathology approaches also fail to capture individual vessel morphology and spatial amyloid distribution across whole-slide images (WSIs). To address this gap, we developed a deep learning framework for reproducible, quantitative analysis of CAA in WSIs.

**Methods:** We analyzed 20 postmortem brain tissue sections from the frontal (n = 10) and occipital cortices (n = 10) of 10 individuals with Alzheimer’s disease pathology obtained from the University of Pittsburgh Alzheimer’s Disease Research Center, which served as the internal development cohort. An independent external cohort consisted of 10 sections (5 frontal and 5 occipital samples) from 5 individuals obtained from the University of Kentucky Alzheimer’s Disease Research Center. We trained and compared three semantic segmentation architectures, a standard U-Net, a dual-attention residual U-Net (DA-ResUNet), and a Swin Transformer-based U-Net (Swin-UNet), using the internal development cohort with slide-level five-fold cross-validation. All models were evaluated on the independent external cohort to assess generalization under domain shift. Based on segmentation performance and computational efficiency, we selected one architecture to generate whole-slide composite segmentation masks for vessel walls, amyloid deposits, and tissue compartments. These masks were subsequently used for deterministic vessel detection, morphometric measurements, and quantification of vascular and perivascular amyloid features through post-processing analysis.

**Results:** All three architectures achieved high segmentation accuracy on the internal cohort, with Dice scores above 90% across vessel walls, amyloid deposits, gray matter, and leptomeninges. The Swin-UNet showed marginally higher performance for vessel segmentation, whereas the DA-ResUNet provided more balanced accuracy and computational efficiency and was selected for downstream analysis. External cohort evaluation demonstrated robust generalization, with attention-enhanced models outperforming the standard U-Net under domain shift. Using the selected model, the pipeline reliably detected valid vessels, excluded non-vascular artifacts, and enabled deterministic extraction of vessel morphometry, vascular and perivascular amyloid burden, and identification of circumferential CAA involvement at the vessel level.

**Discussion:** This framework provides a scalable, interpretable solution for vessel-level CAA analysis, supporting robust geometric and spatial characterization of cerebrovascular pathology and enabling future integration with clinical and genetic studies. Beyond CAA, the modular design allows extension to other vascular pathologies, including arteriolosclerosis, in WSIs, facilitating broader investigation of cerebrovascular disease mechanisms.

## 1 Introduction

Cerebral amyloid angiopathy (CAA) is a cerebrovascular disorder defined by pathological amyloid-beta (Aβ) deposition in the wall of parenchymal and leptomeningeal blood vessels (Revesz et al., 2003). This vascular pathology contributes to cognitive impairment and is a frequent driver of cerebral microhemorrhages and lobar hemorrhages in older adults (Rus Prelog et al., 2025). CAA commonly coexists with Alzheimer’s disease neuropathologic change (ADNC), with autopsy studies showing vascular Aβ in more than 80% of Alzheimer’s disease (AD) brains (Ellis et al., 1996). This vascular pathology adds to the overall disease burden of AD (Ringman et al., 2014).

Despite its clinical and pathological importance, current semiquantitative grading systems limit understanding of how vessel-level features, including Aβ burden, vessel caliber, and anatomic distribution, relate to disease severity and clinical outcomes (Attems et al., 2011; Greenberg et al., 2020). The Vonsattel grading system scores Aβ severity at the level of individual vessels based on wall involvement and structural damage, but it does not capture overall disease burden or anatomic distribution (Vonsattel et al., 1991). The Olichney scoring system, by contrast, summarizes regional involvement but cannot represent mixed patterns of leptomeningeal and parenchymal pathology within the same specimen (Olichney et al., 1995). Both approaches are prone to interobserver variability, and impractical for large-scale quantitative analysis.

Automated analysis of whole slide images (WSIs) using deep learning (DL) offers a path toward objective, scalable, and reproducible quantification of CAA. In principle, such approaches can enable vessel-level morphometric measurements and precise spatial mapping of Aβ burden across large tissue sections (Timakova et al., 2023). However, existing computational methods for CAA analysis remain incomplete. Prior DL frameworks quantify Aβ and vascular features using area-based or density-based measures within manually defined cortical and leptomeningeal regions (Perosa et al., 2021). These approaches do not treat vessels as discrete structural units and thus do not support vessel-level morphometric analysis or assessment of heterogeneity among vessels within the same region.

More recent computational frameworks have moved toward vessel-centered analysis by segmenting individual vessels and deriving morphometric indices, such as wall thickness, in the setting of brain arteriolosclerosis and renal arteriosclerosis (Jin et al., 2024; Lou et al., 2025; Neltner et al., 2014). While these methods introduce precise geometric modeling (e.g., radial sampling), they are optimized for non-Aβ pathologies and often operate on isolated image tiles. As a result, they are ineffective at quantifying the co-localization of Aβ deposition within vessel walls, and they lack the slide-level spatial context required to distinguish vascular compartments, key requirements for the comprehensive assessment of CAA.

Furthermore, in prior works, the choice of DL model architecture has been a key limitation. Single multi-class segmentation networks often suffer from inter-class competition in the presence of severe class imbalance, leading to reduced sensitivity for small or sparse pathological targets such as vascular Aβ deposits (Kervadec et al., 2021). Generalizability also remains a major concern. Many pipelines are developed and evaluated using data from a single institution and are sensitive to variability in staining, tissue processing, and scanning protocols, which restricts their applicability across cohorts and centers (Khened et al., 2021). As a result, reproducible vessel-level quantification and spatial characterization of CAA remain largely unavailable.

In this study, we present a DL pipeline for quantitative analysis of CAA in WSIs. We developed and evaluated two custom segmentation architectures, one transformer-based and one convolutional neural network (CNN)-based and compared their performance with a previously described standard U-Net architecture (Ronneberger et al., 2015) across four targets: Aβ deposits, vessel walls, leptomeningeal space (LS), and gray matter (GM). We trained independent binary models for each target and fused their outputs into a composite semantic map. Using data from two institutions, we assessed robustness across staining and scanning conditions. Finally, we utilized these maps to spatially assign individual vessels to anatomical compartments and derive vessel-level morphometric and Aβ burden measurements. This framework enables reproducible, vessel-level quantification of CAA beyond conventional semiquantitative grading systems.

## 2 Materials

### 2.1 Cohort selection and tissue acquisition

We obtained postmortem human brain tissue sections from 15 individuals with autopsy confirmed AD through the Alzheimer’s Disease Research Centers (ADRC) at the University of Pittsburgh (Pitt) (n = 10; internal training and validation data set) and the University of Kentucky (UKY) (n = 5; independent external test data set). From each individual, we analyzed two cortical tissue sections, one from the frontal cortex and one from the occipital cortex. Demographic characteristics and neuropathologic assessments for the Pitt and UKY cohorts are summarized in Table 1 and Table 2, respectively. Informed consent for autopsy and the research use of tissue was obtained from next-of-kin of all brain donors. The study was approved by the Committee for Oversight of Research and Clinical Training Involving Decedents (CORID) at Pitt and the UKY Institutional Review Board. All data were de-identified before analysis in accordance with institutional policies and applicable privacy regulations.

**Table 1.**
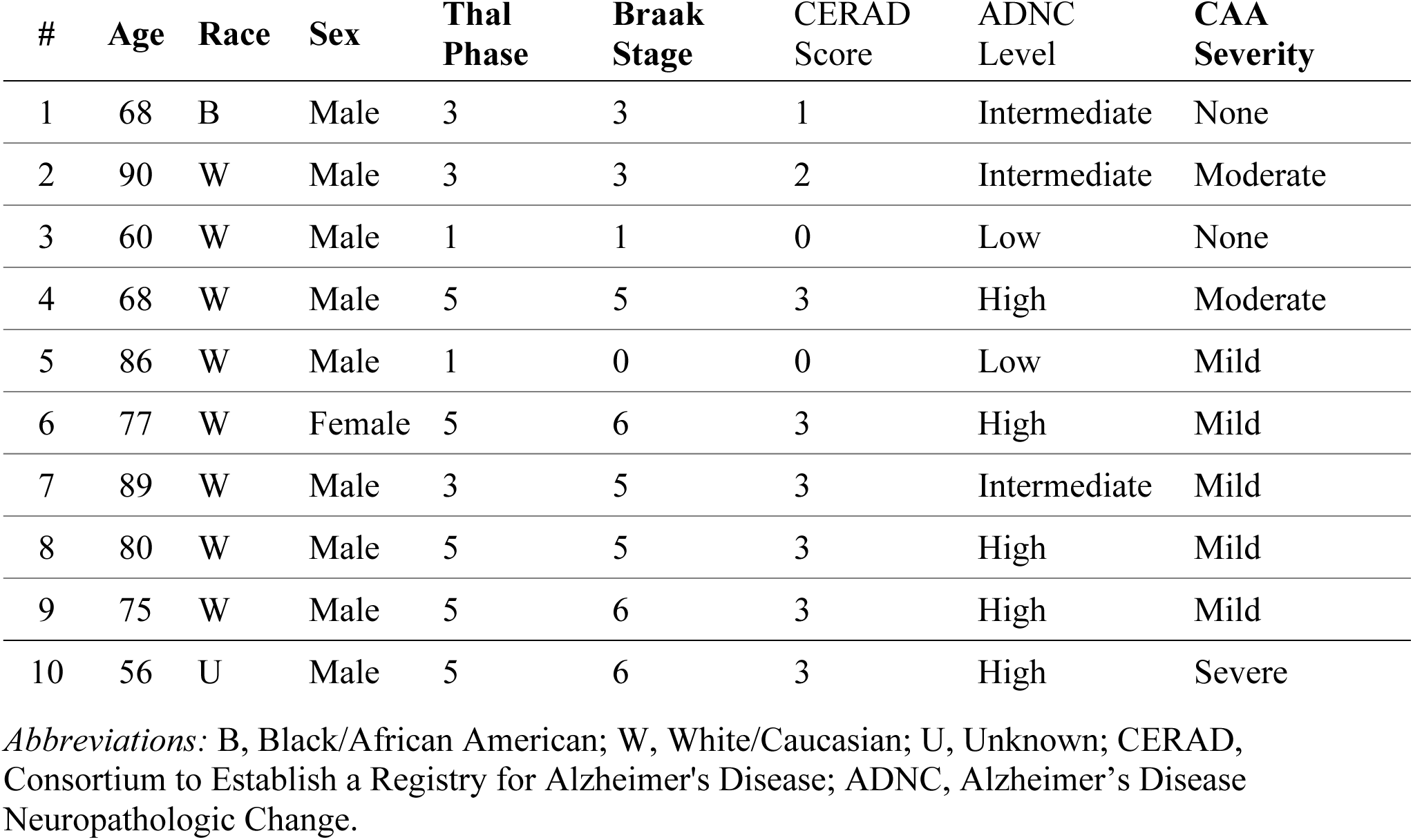
Demographic and neuropathologic characteristics of the Pitt cohort.

**Table 2.**
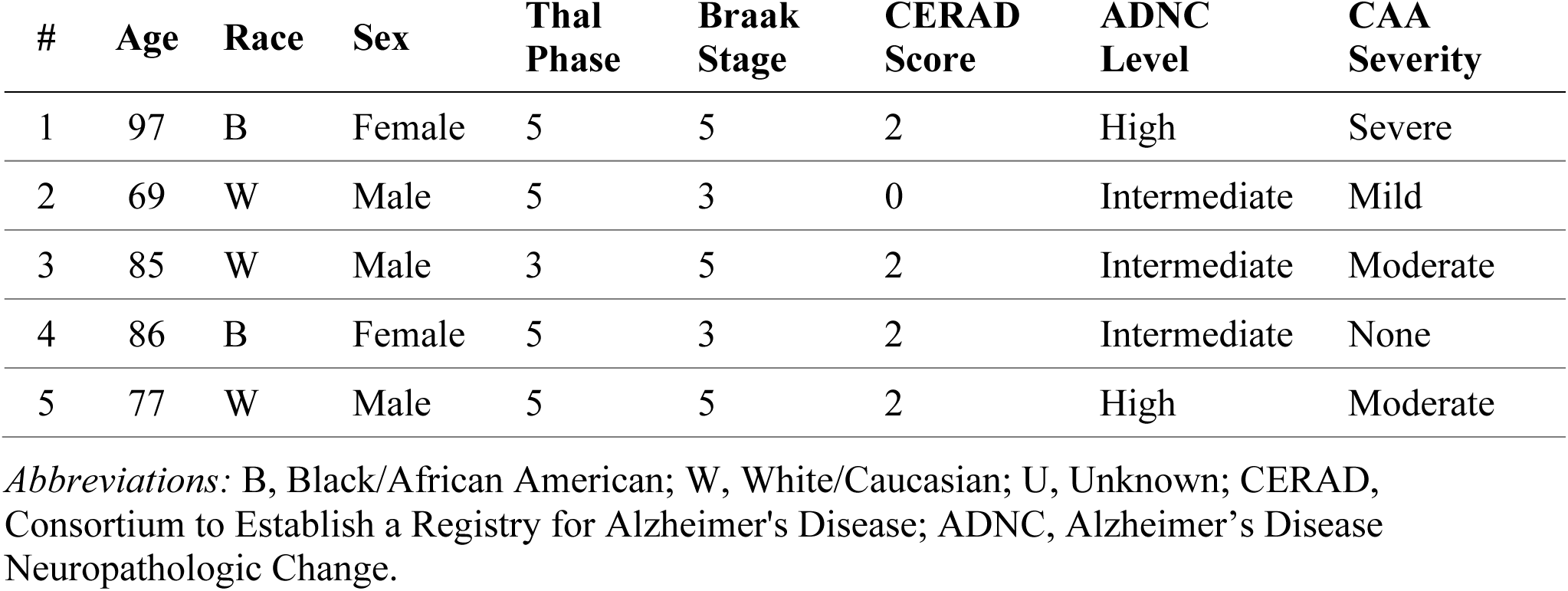
Demographic and neuropathologic characteristics of the UKY cohort.

Diagnosis of presence and severity of ADNC was based on established criteria (Montine et al., 2012), incorporating Braak neurofibrillary tangle (NFT) staging (Braak et al., 2006), Thal Aβ phases (Thal et al., 2002), Consortium to Establish a Registry for Alzheimer’s Disease (CERAD) neuritic plaque scores (Mirra et al., 1991), and overall ABC scores, as performed by a board-certified neuropathologist. We sampled two cortical regions from the left cerebral hemisphere of each case: the dorsolateral prefrontal cortex at the level of the midfrontal gyrus and the medial occipital cortex including the primary visual cortex. This strategy yielded a total dataset of 30 tissue sections. Each section comprised the full-thickness cortex, capturing cortical GM and underlying white matter (WM).

### 2.2 Tissue processing

Sections from the Pitt cohort were cut at a thickness of 5 µm from formalin-fixed, paraffin-embedded (FFPE) blocks. Slides were first deparaffinized, rehydrated, and treated with methanol hydrogen peroxide to block endogenous peroxidase activity, followed by pretreatment with 90% formic for 40 min. Aβ immunohistochemistry was performed using antibody NAB228 (1:4000; Cell Signaling Technology, Danvers, MA) for 45 min at room temperature. Signal detection was carried out using a biotinylated mouse IgG secondary antibody (1:200 for 30 minutes), the ABC Elite detection system, and Nova Red chromogen. Slides were then counterstained with hematoxylin. No vessel-specific immunohistochemical markers were applied.

At UKY, slides were cut from FFPE blocks at 8 µm thickness and were also immunostained using the NAB228 anti-Aβ antibody (1:2000, with formic acid pretreatment, with 3,3’-diaminobenzidine chromogen). As a result, staining procedures were similar but not identical across institutions.

### 2.3 Image digitization

The glass slides were digitized at Pitt using the Leica Aperio AT2 scanner at 40X nominal magnification. The scanner metadata indicated a spatial resolution of approximately 0.25 μm per pixel. We received slides from the UKY as WSIs, also scanned at 40X magnification using an AT2 scanner at UKY and shared through access to the UKY Digital Slide Archive. All slides were stored in SVS format using JPEG compression.

### 2.4 Computational environment

This study utilized computing resources provided by the Center for Research Computing and Data at Pitt. All model training and inference was performed on a single NVIDIA A100 GPU. The computational pipeline was implemented in Python using the PyTorch framework. Detailed software specifications and library versions are listed in Table 3. To improve computational efficiency, we employed mixed-precision training (Dörrich et al., 2023). We also fixed random seeds across all libraries to ensure reproducibility.

**Table 3.**
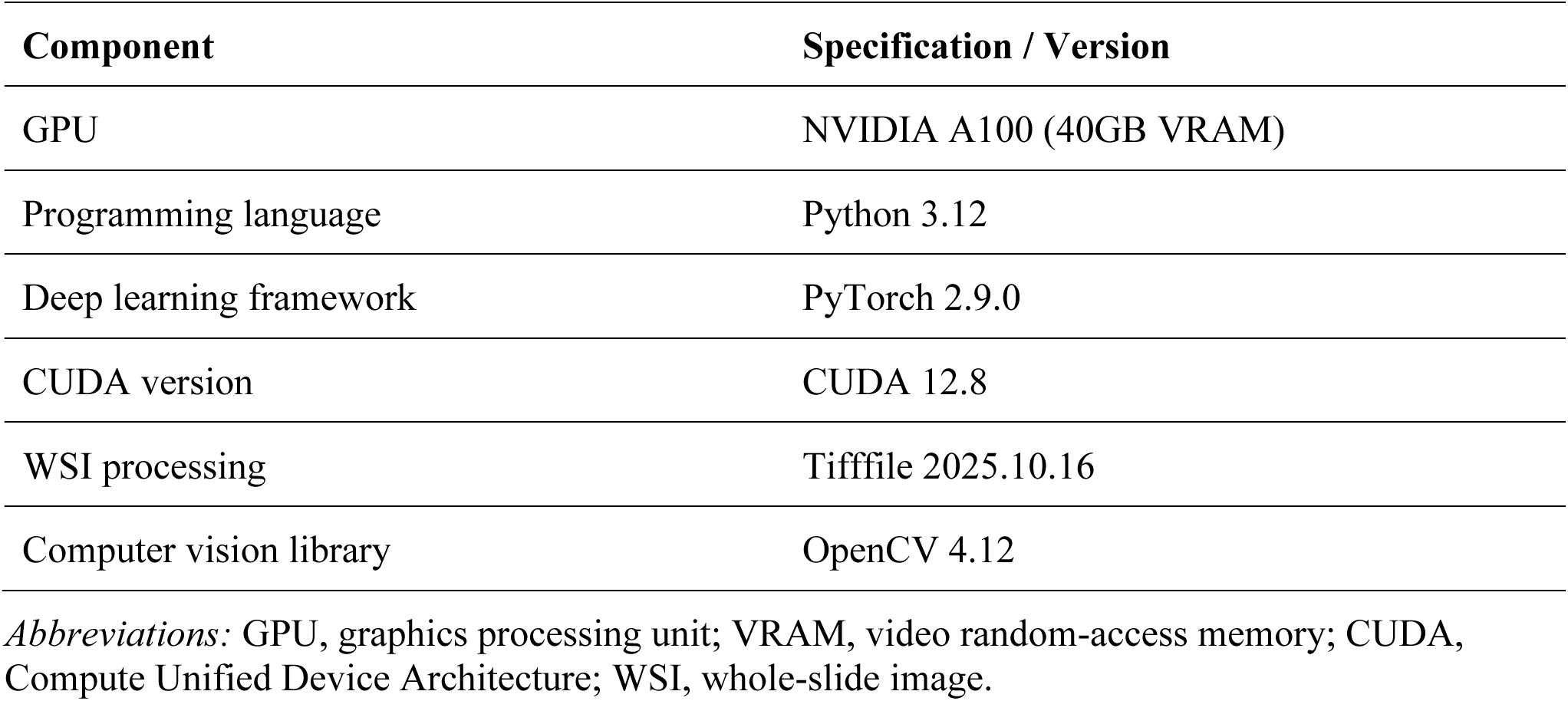
Computational environment and software specifications.

## 3 Methods

### 3.1 Annotation strategy

All reference annotations were generated using QuPath version 5.1 (Bankhead et al., 2017). A single annotator (H.T.) manually delineated regions of interest under the direct supervision of a board-certified neuropathologist (J.K.). The annotator produced binary pixel-level masks for four distinct target classes: Aβ deposits, vessel walls, GM, and LS (Figure 1). To ensure comprehensive segmentation, Aβ labels encompassed both vascular and parenchymal deposits, while vessel wall labels included both unaffected and Aβ-laden vessels. These finalized masks served as the ground truth for model training, validation, and testing.

**Figure 1.**
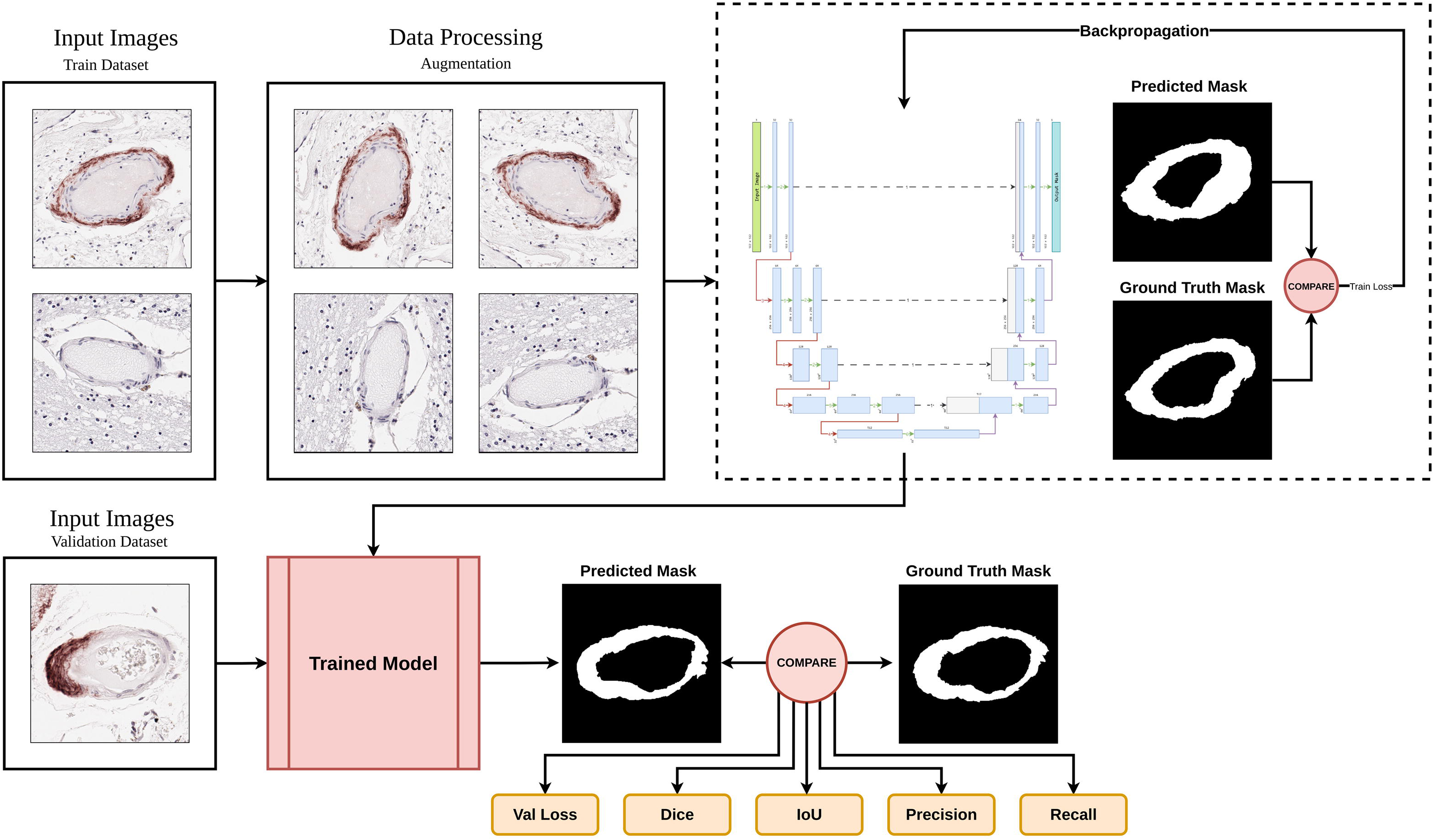
Annotation strategy and ground truth generation. Representative examples of the annotation strategy used to create the four training, validation, and testing datasets. Columns show the raw input image (left column), the expert annotation overlay highlighted in yellow (middle column), and the final binary ground truth mask (right column) with the white color indicating the target object. To capture appropriate structural detail, vessel wall and Aβ were sampled at 40X magnification, while leptomeningeal space and gray matter were sampled at 10X.

### 3.2 Tile extraction

For Aβ and vessel wall segmentation tasks, we extracted image tiles at 40X magnification with tile dimensions 512 x 512 pixels and no overlap, as these targets require high spatial resolution to capture fine structural detail and sparse pathology. For GM and LS segmentation, we extracted tiles at 10X magnification using the same pixel dimensions, as these tissue compartments span large contiguous regions. We excluded tiles with minimal tissue content by applying a tissue coverage threshold of 5 percent. Tiles below this threshold were removed prior to model training, validation, and testing.

### 3.3 Data partitioning

#### 3.3.1 Model training and validation

We split the dataset by individual WSIs to ensure the training and validation sets remained strictly independent. Digitized slides from the Pitt cohort were used for model development. This set included 10 frontal cortex sections and 10 occipital cortex sections. These slides were used exclusively for training and validation. We implemented a five-fold cross-validation scheme on this development set. Crucially, all tiles derived from a single WSI were assigned to the same partition within each fold to prevent data leakage between training and validation subsets (Allgaier & Pryss, 2024). An overview of this workflow is shown in Figure 2. Table 4 provides a detailed summary of the dataset composition and the total number of tiles extracted for each target class.

**Figure 2.**
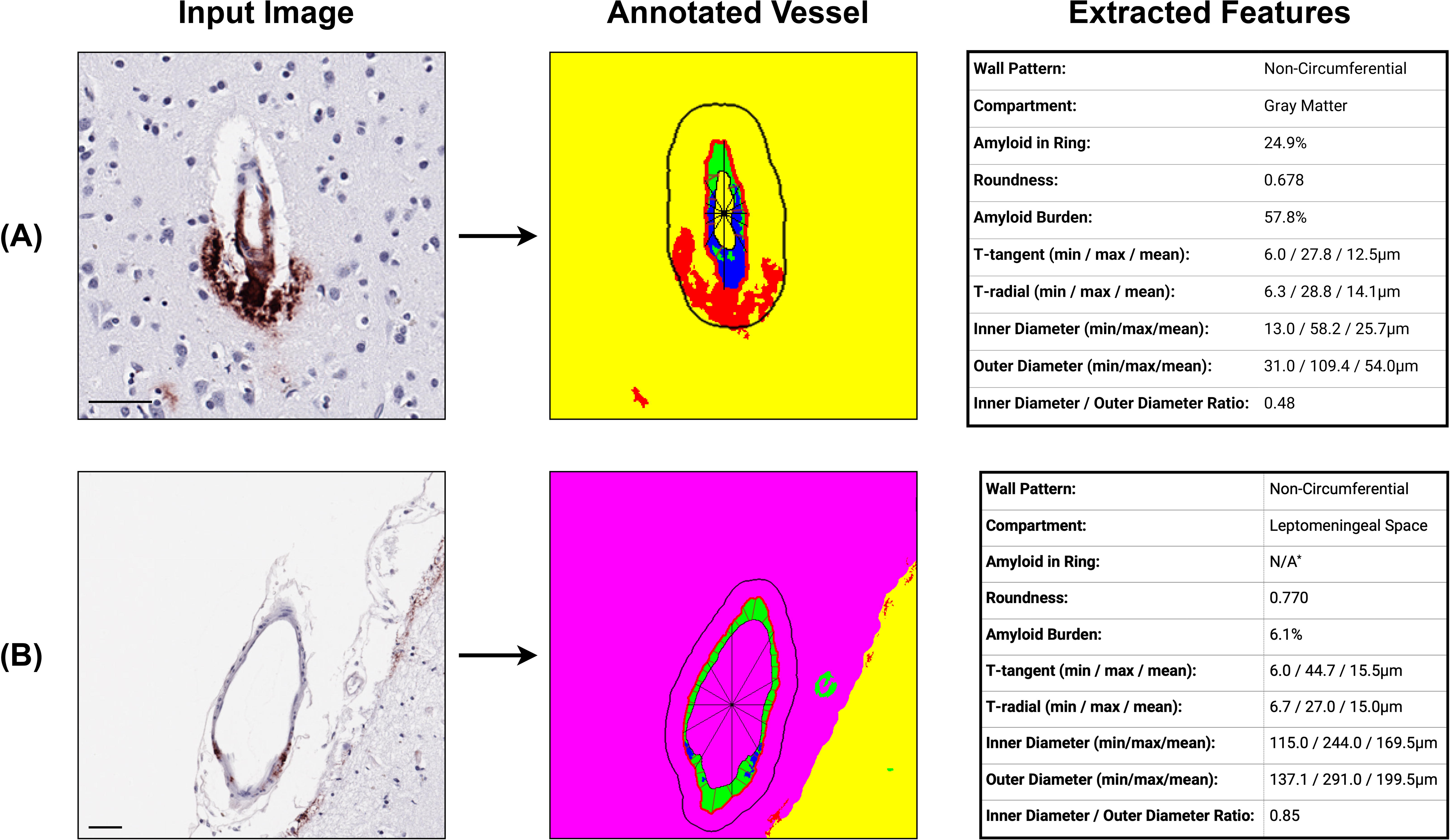
Overview of the training and validation workflow used for semantic segmentation. The top panel shows the training phase, in which input images from the training dataset undergo data augmentation before being processed by the segmentation model. Predicted masks are compared with ground truth annotations to compute the training loss, which is used to update the network through backpropagation. The bottom panel shows the validation phase, in which unseen input images are passed through the trained model. The resulting predicted masks are compared with ground truth masks to evaluate model performance using validation loss, Dice coefficient, intersection over union, precision, and recall.

**Table 4.**
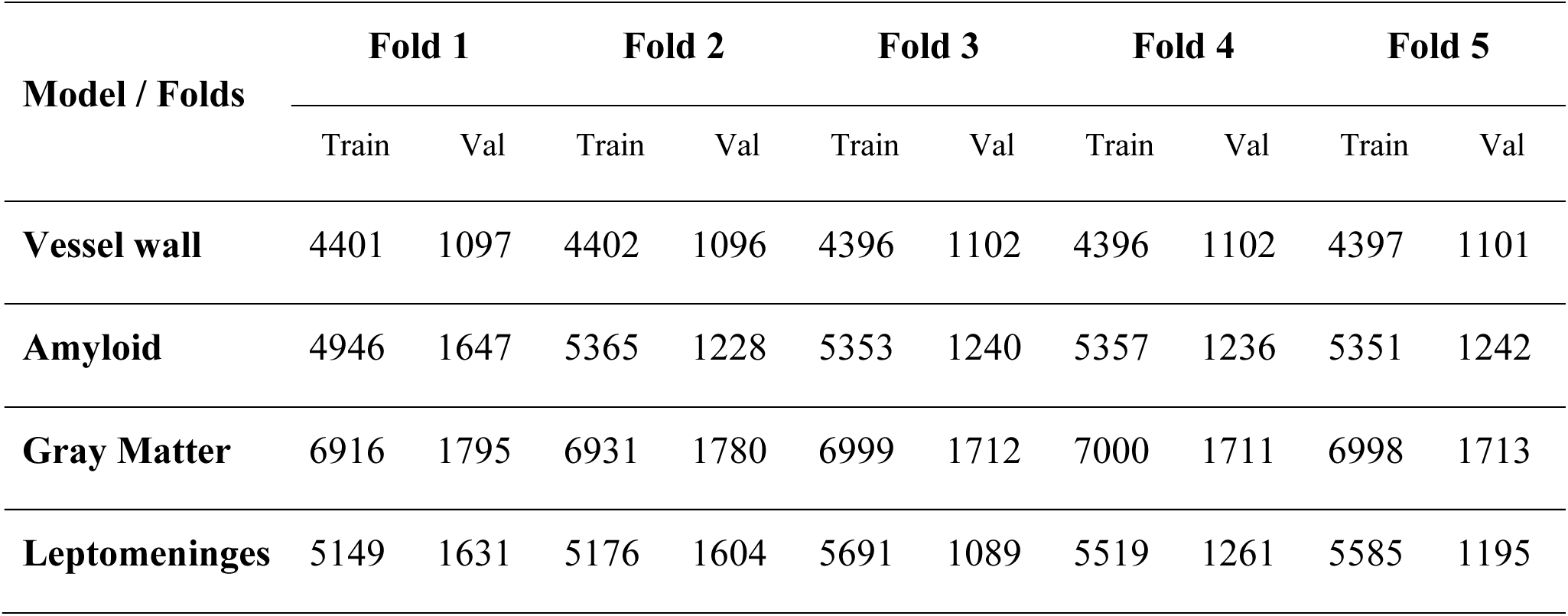
Number of training (Train) and validation (Val) tiles per fold for each model across five-fold cross-validation setup.

#### 3.3.2 Model testing

To evaluate generalization to independent datasets, we performed external cohort testing using WSIs from the UKY. These data were strictly held out during all phases of model development, including training, cross-validation, and hyperparameter optimization. We extracted 500 tiles per target class (vessel walls, Aβ deposits, GM, and LS) from 10 WSIs (5 frontal cortex and 5 occipital cortex), using the same magnification settings as in the development phase. Ground truth masks were generated by the same annotator. We applied the optimal checkpoints from the internal development set directly to these external tiles without any additional fine-tuning or domain adaptation.

### 3.4 Data augmentation

During training, we applied paired geometric transformations to input images and their corresponding ground truth masks to preserve spatial correspondence. These transformations were limited to random horizontal and vertical flips (Shorten & Khoshgoftaar, 2019). Following geometric transformations and intensity scaling, we applied image-only appearance augmentation with a probability of 0.5. This augmentation perturbed stain appearance in optical density space to simulate staining and scanner-related variability and included Gaussian blur and additive noise (Tellez et al., 2019). We did not apply any data augmentation during validation or testing. Figure 3 shows examples of data augmentation used during model training

**Figure 3.**
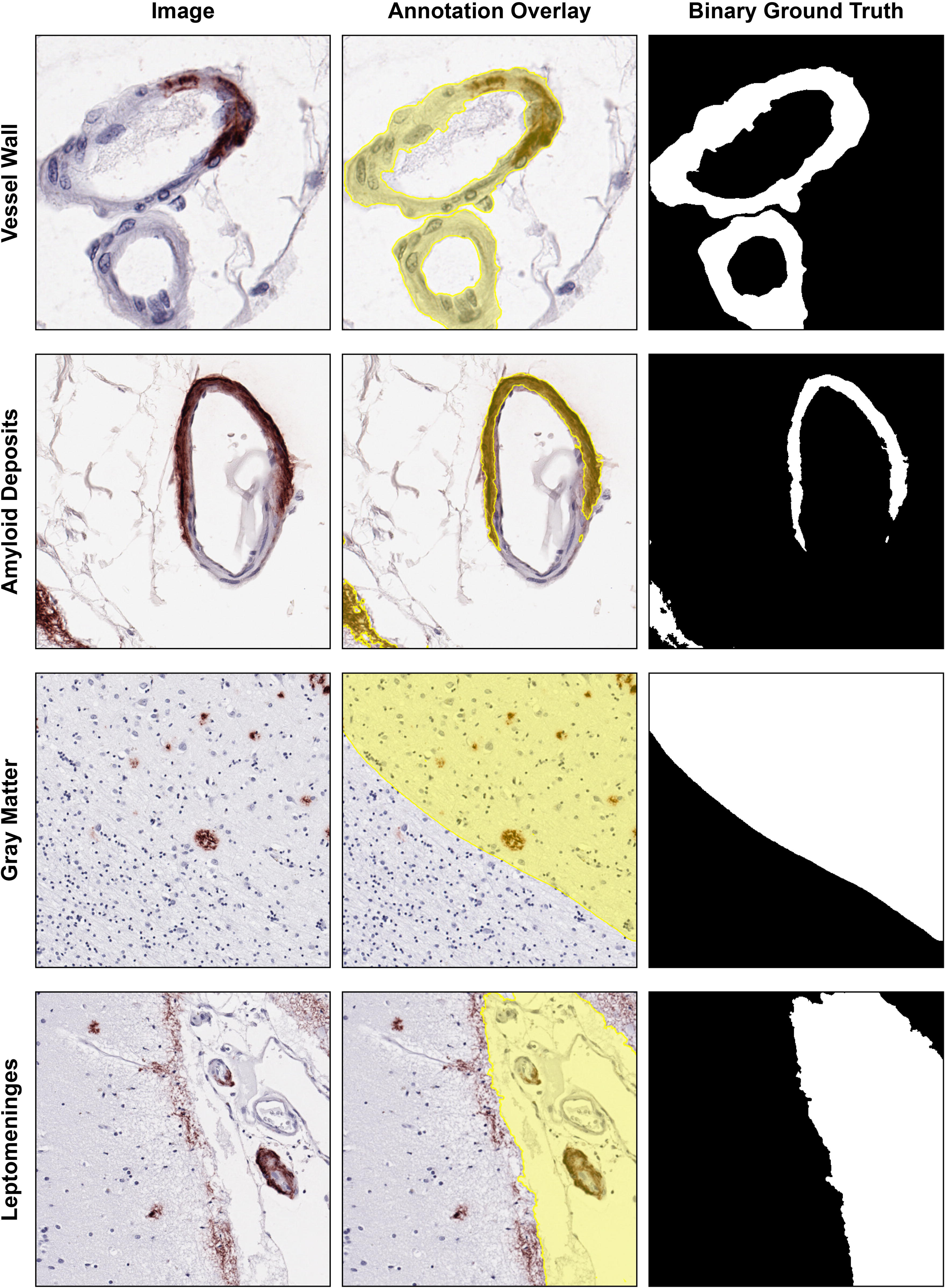
Examples of data augmentation used during model training. The Baseline column shows the original image. The Geometric augmentation column shows paired spatial transformations applied to images and masks, including rotation and horizontal and vertical flips. The Stain augmentation column illustrates color and intensity perturbations applied in optical-density space to model variability in hematoxylin and DAB staining. The Scanner augmentation column shows photometric variations, including channel mixing, intensity scaling, gamma adjustment, blur, and additive noise, used to simulate scanner-related variability. Scale bar = 100μm.

### 3.5 Model architectures

We developed three U-Net-style semantic segmentation architectures. Each model mapped an RGB (red, green, and blue) tile x∈R^3×H×W^ to a single channel logit map z∈R^1×H×W^. A logit map is the raw output of the network before conversion to probabilities. We converted these logits to pixel-wise probabilities using a sigmoid function, y"=σ(z), which maps each pixel value to a range between 0 and 1 and allows the model to estimate the probability that each pixel belongs to the target class (Hao et al., 2020). Table 5 details the computational specifications for each architecture, including the total number of trainable parameters and floating-point operations (FLOPs), which estimate the amount of computation required to process one input tile.

**Table 5.**
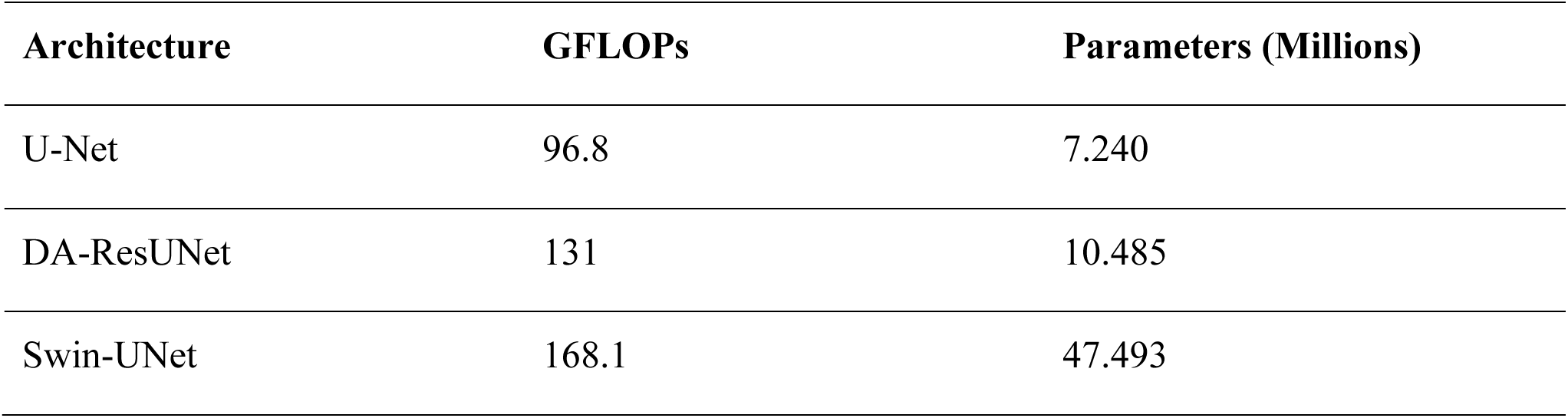
Computational complexity and model size of the implemented architectures.

#### 3.5.1 Standard U-Net

We implemented a standard U-Net architecture following the original design (Ronneberger et al., 2015). The network consisted of a symmetric encoder-decoder structure with skip connections between corresponding resolution levels (Figure 4), in which the encoder extracts features, and the decoder reconstructs the segmentation mask while the skip connections preserve fine spatial detail.

**Figure 4.**
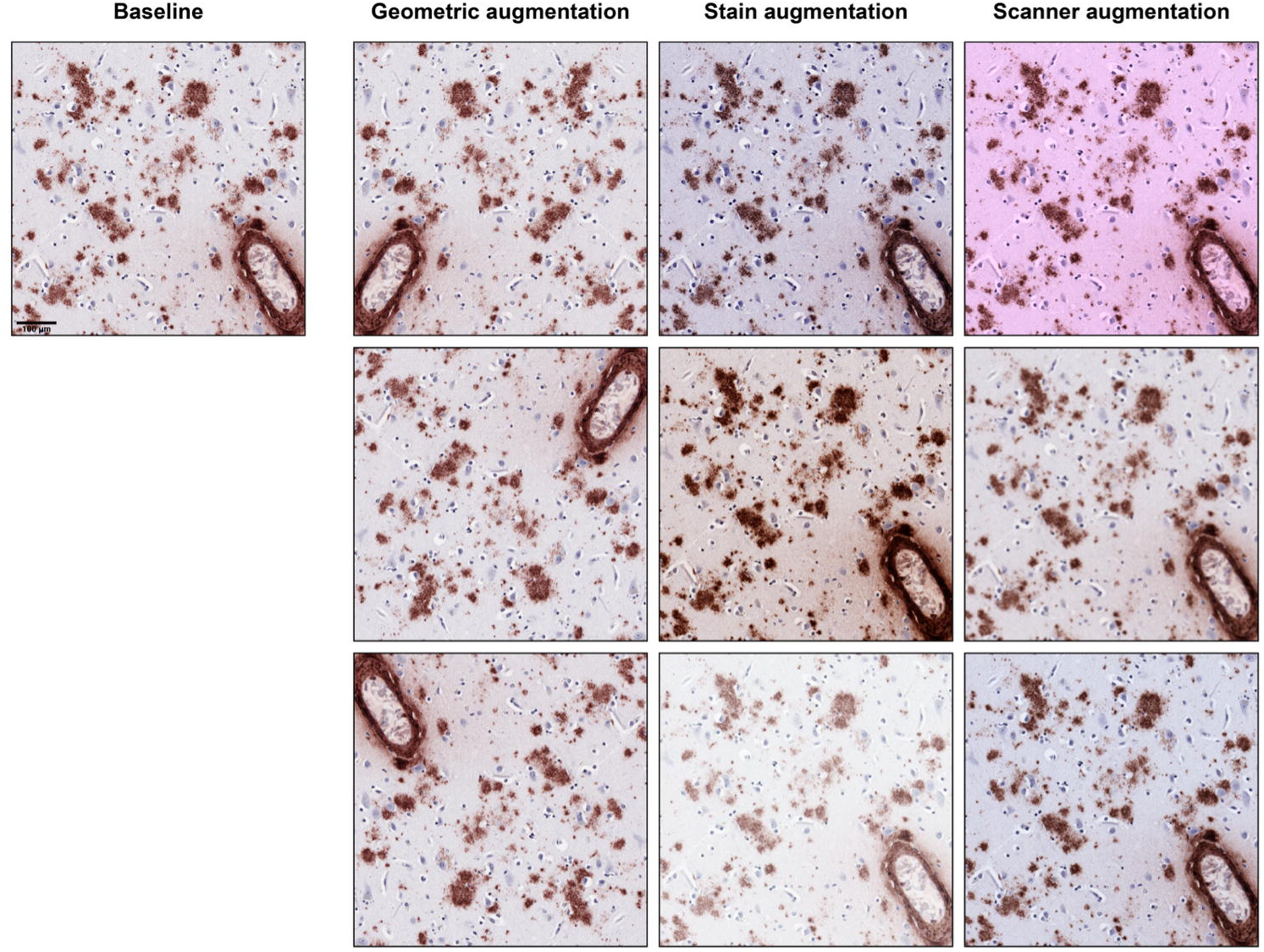
Architecture of the standard U-Net. Overview of the standard U-Net encoder-decoder architecture with skip connections.

##### 3.5.1.1 Encoder

The encoder included four stages with progressive spatial downsampling. Downsampling reduces the spatial size of the feature maps while increasing the effective receptive field, which allows the model to use a larger image context. Each stage applied two consecutive 3 x 3 convolutional layers followed by batch normalization and Rectified Linear Unit (ReLU) activation. Convolutional layers learn local image patterns such as edges, textures, and shapes. Batch normalization stabilizes training by standardizing feature values. ReLU is a nonlinear activation function that sets negative values to zero and helps the network learn more complex patterns. Downsampling between stages was performed using max pooling. The number of feature channels increased with depth, starting from 32 channels in the first stage.

##### 3.5.1.2 Decoder

The decoder mirrored the encoder and progressively restored spatial resolution. At each stage, feature maps were upsampled and concatenated with the corresponding encoder features. The fused features were refined using two consecutive convolutional layers with batch normalization and ReLU activation. A final 1 x 1 convolution produced a single-channel logit map for segmentation.

#### 3.5.2 Dual-attention residual U-Net

The first model extended the standard U-Net architecture (Ronneberger et al., 2015) by incorporating residual learning (He et al., 2016) and attention mechanisms to enhance feature representation while preserving spatial detail (Woo et al., 2018). Residual learning uses shortcut connections to help information and gradients pass more easily through the network during training. Figure 5 shows an overview of this model architecture.

**Figure 5.**
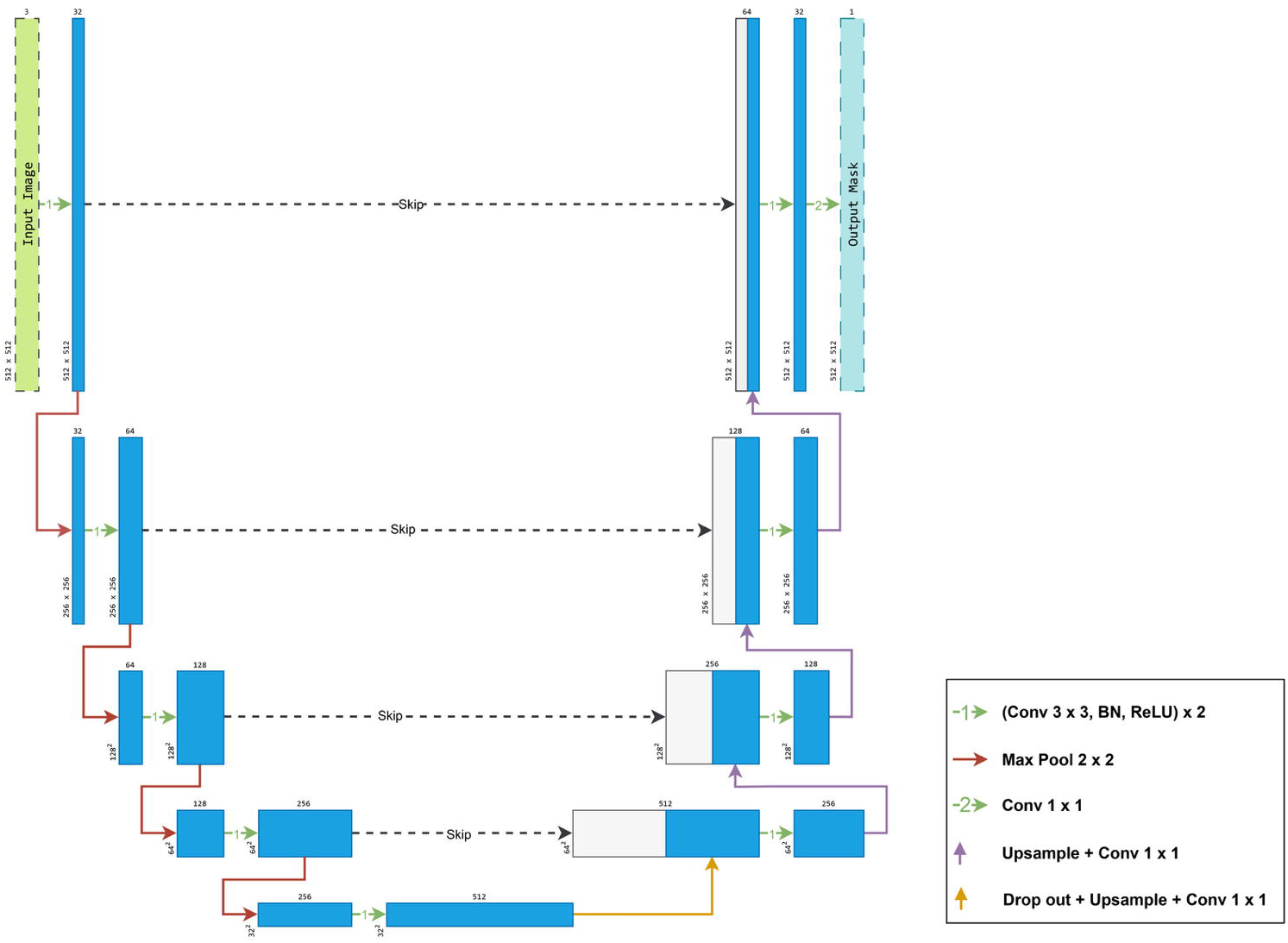
Architecture of the dual-attention residual U-Net (DA-ResUNet). Overview of the U-Net-based encoder-decoder architecture with skip connections and integrated attention and residual components used for semantic segmentation.

##### 3.5.2.1 Encoder

The encoder consisted of four stages with progressive spatial downsampling. We designed a hybrid block strategy to optimize gradient flow at different scales. In the shallow encoder stages (stages 1 and 2), we applied three consecutive convolutional blocks, each composed of a 3 x 3 convolution, batch normalization, and Gaussian Error Linear Unit (GELU) activation (Purwanti et al., 2025). We used GELU because it is a smoother activation function than ReLU and often supports stable optimization in modern architectures. In the deeper encoder stages (stages 3 and 4), we replaced standard convolutional blocks with post-activation residual blocks to facilitate gradient propagation and stabilize training at higher semantic levels (He et al., 2016) (Figure 6). Inspired by convolutional block attention module (CBAM)-Residual U-Net, we applied a CBAM after each encoder stage and before downsampling (Shah & Kang, 2023). CBAM is a lightweight attention module that sequentially applies channel attention and spatial attention. Channel attention helps the model weigh which feature channels are most informative, whereas spatial attention helps the model emphasize where the important signal is located within the image. The module applied channel attention followed by spatial attention (Woo et al., 2018).

**Figure 6.**
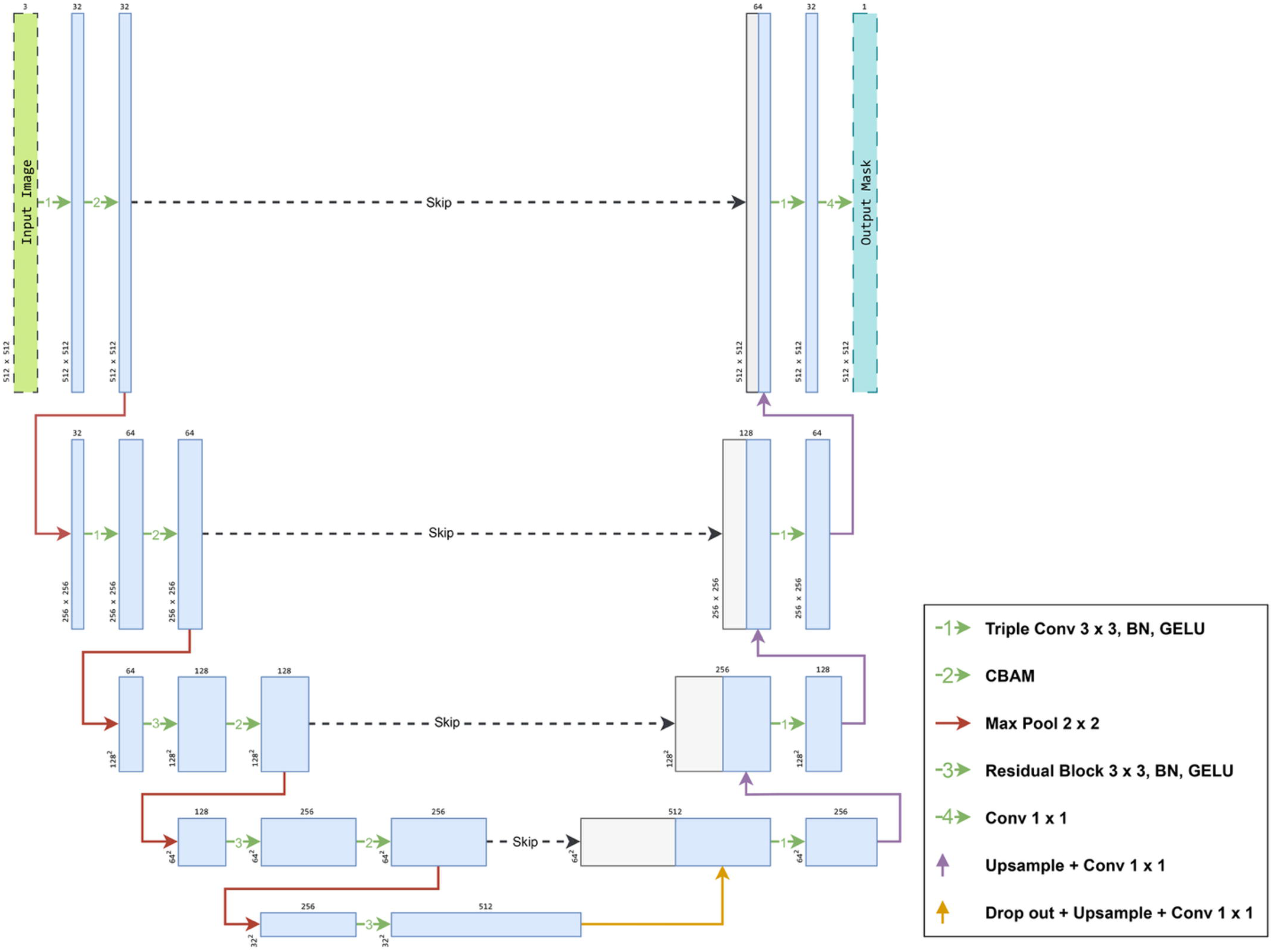
Post-activation residual block architecture. The block consists of two sequential 3 x 3convolutional layers, each followed by normalization, with GELU activation applied after the residual addition. A skip connection adds the input feature map to the transformed features to facilitate gradient flow. In the Swin-UNet model, normalization corresponds to instance normalization, whereas in the dual-attention residual U-Net, normalization corresponds to batch normalization.

##### 3.5.2.2 Decoder

The decoder progressively recovered spatial dimensions using an upsampling scheme designed to mitigate the checkerboard artifacts often associated with transposed convolutions (Kinoshita & Kiya, 2020). At each decoding stage, we applied bilinear interpolation with a scale factor of two, followed by a 1 x 1 convolution to reduce channel dimensionality (Zhang et al., 2019). We concatenated the upsampled features with the corresponding encoder features through skip connections. The fused features were refined using three consecutive 3 x 3 convolutions, batch normalization, and GELU layers. A final 1 x 1 convolution mapped the decoder features to a single-channel logit map.

#### 3.5.3 Hybrid Swin transformer U-Net

The second model replaced the convolutional encoder with a hierarchical transformer encoder based on Swin Transformer V2 (Liu et al., 2022). Inspired by SwinUNETR-V2, convolutional layers were integrated into the architecture to reinforce local spatial coherence and stabilize training (He et al., 2023). Figure 7 illustrates the architecture of this model.

**Figure 7.**
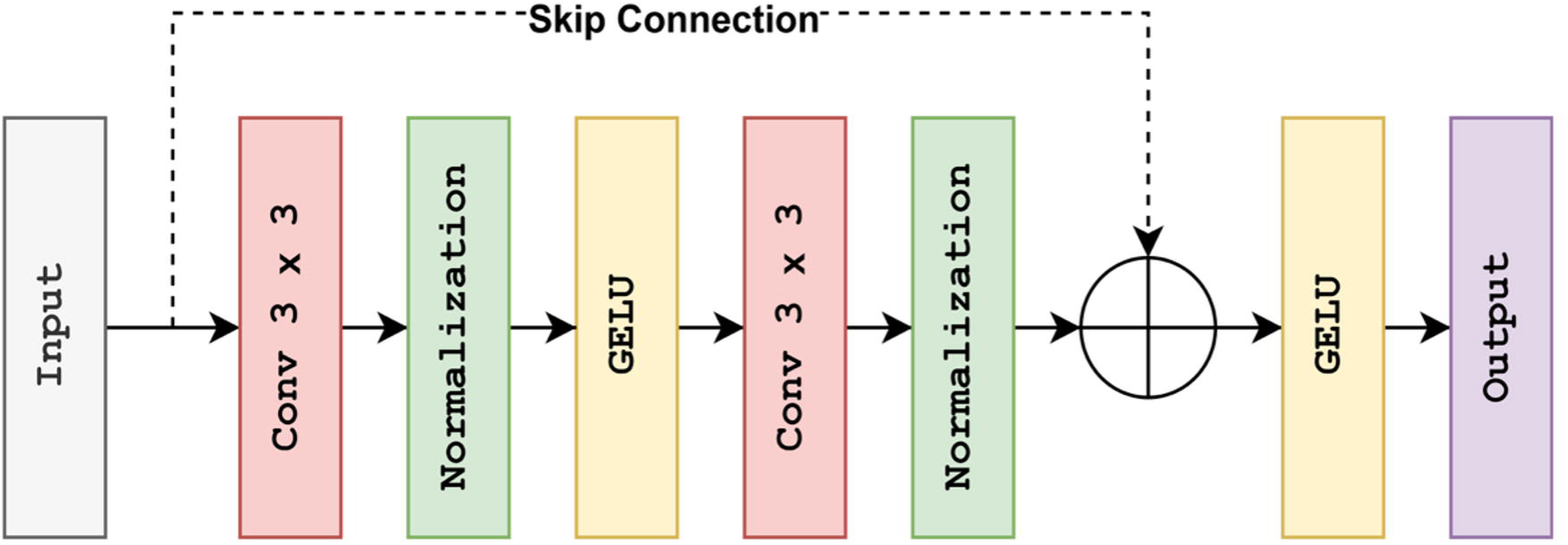
Architecture of the Swin-UNet segmentation model. The network combines a hierarchical Swin Transformer V2 encoder with residual convolutional blocks and a U-Net-style decoder. Patch embedding and successive encoder stages capture multi-scale contextual features, while skip connections and fusion convolutions integrate encoder and decoder representations. The model produces a single-channel segmentation mask through a final 1 x 1 convolutional output head.

##### 3.5.3.1 Patch embedding

The model partitioned each input image into nonoverlapping 3 x 3 patches. This step breaks the image into small local regions so they can be processed as tokens by the transformer. A strided convolution projected these patches into a higher-dimensional embedding space, preserving spatial structure while converting local neighborhoods into token representations.

##### 3.5.3.2 Encoder

Each encoder stage began with a pre-activation residual convolutional block (Figure 8). This block applied instance normalization and GELU activation prior to a 3 x 3 convolution. Instance normalization normalizes each image sample independently, which can be helpful when feature statistics vary across samples. This design stabilized feature statistics and reinforced local spatial structure before attention-based processing (Wu et al., 2021). Following convolutional refinement, features were processed using Swin Transformer V2 blocks with window-based multi-head self-attention. In this setting, self-attention allows the model to compare information across different image locations, while the window-based design limits this operation to local regions to reduce computational cost. Attention alternated between non-shifted and cyclically shifted windows to enable information exchange across window boundaries while maintaining linear computational complexity with respect to image size (Liu et al., 2022).

**Figure 8.**
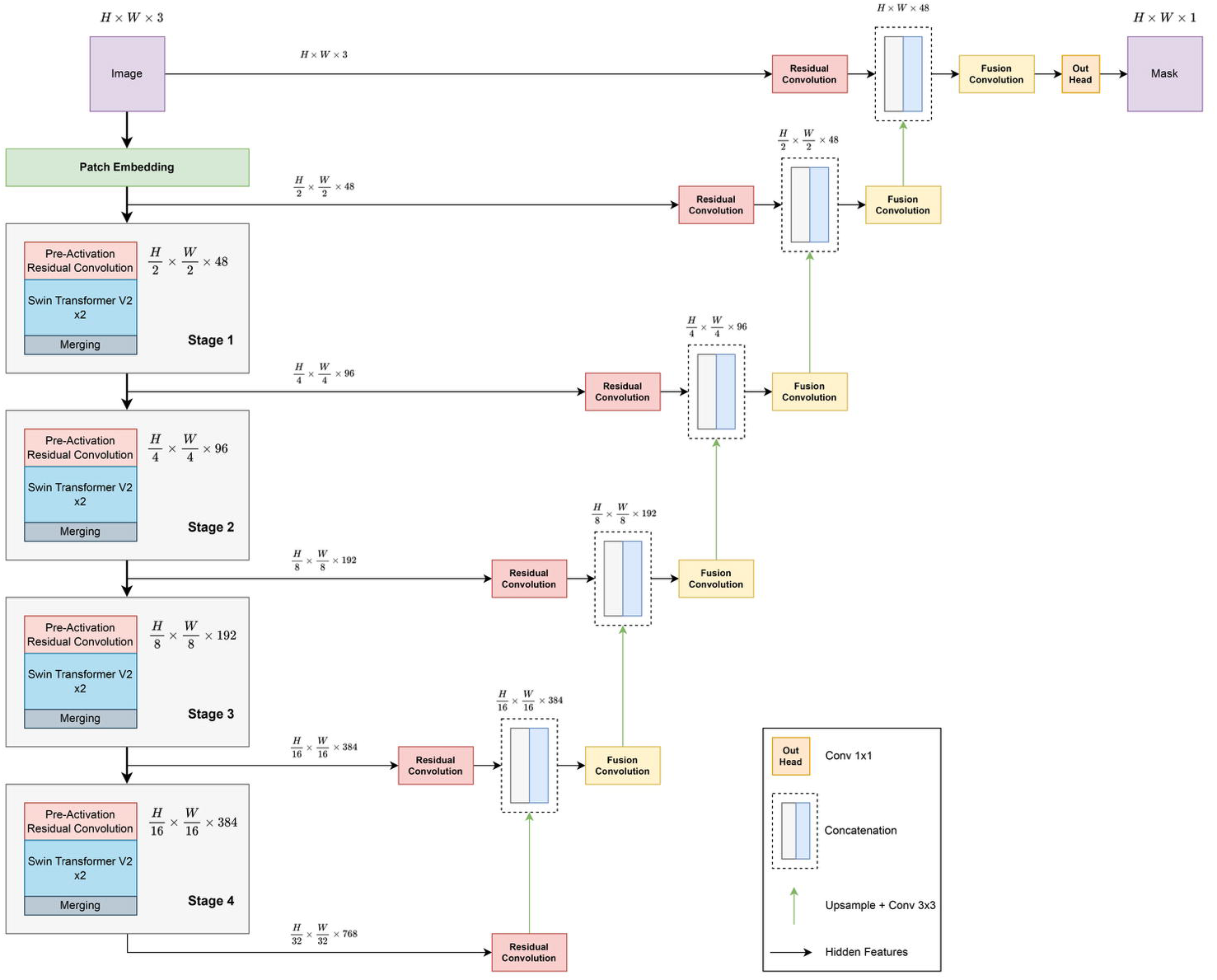
Pre-activation residual block architecture. The block applies normalization and GELU activation before each 3 x 3 convolution, with a skip connection that adds the input feature map to the transformed features. In the dual-attention residual U-Net, normalization corresponds to batch normalization.

Each transformer block followed a residual formulation and included layer normalization and a multilayer perceptron with GELU activation. Stochastic depth was applied to residual paths during training to regularize deep representations. Stochastic depth randomly drops some residual paths during training and acts as a form of regularization. Between encoder stages, the model applied patch merging. This operation concatenated neighboring tokens in a space-to-depth manner and projected them through a linear layer, reducing spatial resolution while increasing channel capacity.

##### 3.5.3.3 Decoder

The decoder mirrored the encoder hierarchy. At each level, bilinear upsampling doubled the spatial resolution, followed by convolution to reduce channel dimensionality. Skip features from the encoder were first processed using residual convolutional blocks to align semantic representations before concatenation. This step helped make the encoder and decoder features more compatible before fusion. The fused features were then refined using additional residual convolutional blocks. A final 1 x 1 convolution produced the binary segmentation logit map.

We computed FLOPs using a standard input tile size of 512 x 512 pixels. Table 5 lists the computational cost and the total number of trainable parameters for the U-Net, Swin-UNet, and DA-ResUNet architectures.

### 3.6 Model training strategy

We fixed random seeds and enabled deterministic computation to improve reproducibility across runs. We trained models for 150 epochs, evaluating performance on the held-out validation fold at the end of each epoch. Table 6 summarizes the hyperparameter settings used for model training.

**Table 6.**
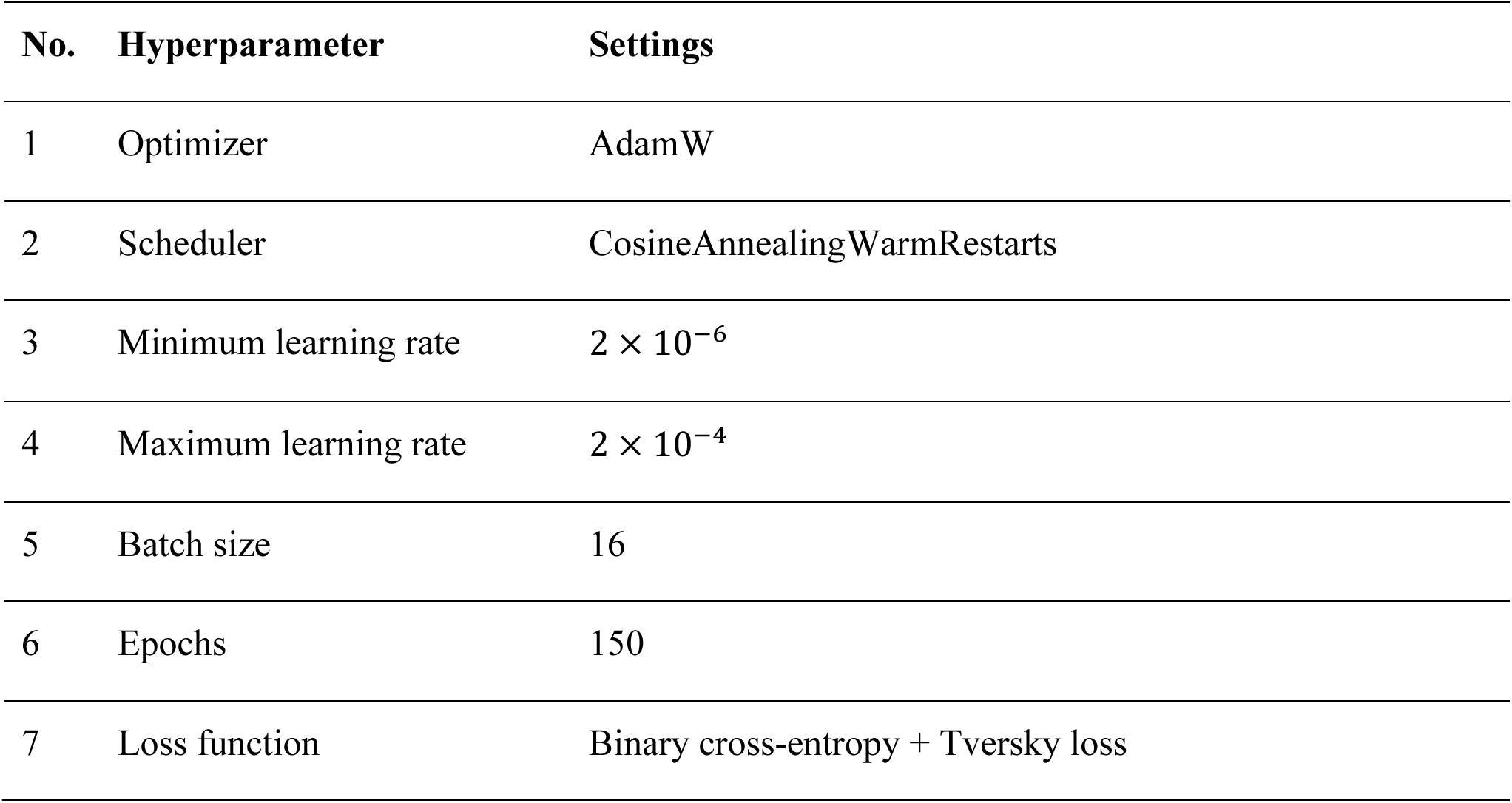
Hyperparameter settings for all models.

We monitored the validation loss to select the optimal model checkpoint. To assess segmentation performance, we calculated the Dice coefficient (F1-score), precision, and recall (Müller et al., 2022). The following equations define the metrics:

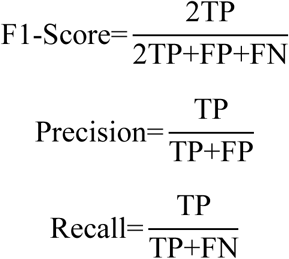

where TP denotes true positives, FP false positives, and FN false negatives.

### 3.7 Loss function and optimization

We trained the models using a composite loss function to address class imbalance and effectively segment structures of varying sizes. We combined Binary Cross-Entropy (L_BCE_) to maximize pixel-level classification accuracy with Tversky Loss (L_Tversky_) to optimize region-level overlap (Taghanaki et al., 2019). We used this combination because Binary Cross-Entropy focuses on individual pixel errors, whereas Tversky Loss places more emphasis on overlap of the segmented region. We defined total loss L_total_ as:

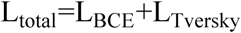

We calculated the L_BCE_, which penalizes probabilistic errors at the individual pixel level, as:

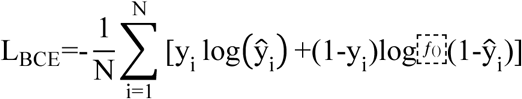

where y_i_ represents the ground truth label (0 or 1) and y"_i_ denotes the predicted probability for pixel i (Janthakal & Hosalli, 2021).

We employed Tversky Loss to improve sensitivity for small or sparse targets by balancing the penalties for FP and FN (Salehi et al., 2017). We defined this loss as:

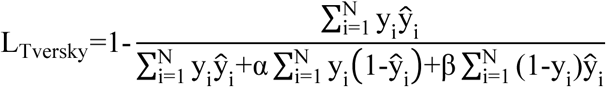

where α and β are hyperparameters that control the trade-off between false negatives and false positives, respectively.

We optimized model parameters using the AdamW optimizer (Loshchilov & Hutter, 2017). To stabilize training and escape local minima, we adjusted the learning rate using a cosine annealing schedule with warm restarts applied at the batch level (Loshchilov & Hutter, 2016). This schedule gradually lowers the learning rate and then periodically resets it, which can help training continue improving instead of becoming stuck too early. We utilized this schedule to promote rapid initial convergence while reducing the risk of premature optimization stagnation.

### 3.8 Multi-resolution inference strategy

We utilized the four trained binary segmentation models targeting vessel walls, Aβ deposits, GM, and LS to process WSIs. To balance segmentation accuracy with computational efficiency, we performed inference at different spatial resolutions of the image pyramid depending on the target structure. GM and LS were segmented at a downsampled pyramid level corresponding to a fourfold reduction in resolution, as these compartments span large contiguous regions and do not require cellular-level detail. In contrast, vessel wall and Aβ segmentation were performed at native image resolution to preserve fine structural detail and to capture sparse Aβ deposits and vessel morphometry. We performed inference using a sliding window approach with tiles of size 2048 x 2048 pixels and an overlap of 1024 pixels. The overlap reduced edge effects at tile boundaries and improved spatial continuity in the reconstructed predictions. To increase robustness at inference time, we applied test-time augmentation (TTA) (Moshkov et al., 2020). For each tile, we generated predictions from the original image and from its horizontally and vertically flipped versions. We averaged these predictions to produce the final probability map for each tile.

### 3.9 Composite mask construction

We synthesized a single composite segmentation map by fusing the outputs from the four independent binary models using a bitwise encoding strategy. We assigned a unique power-of-two label to each model prediction: LS (2^0^=1), GM (2^1^=2), vessel wall (2^2^=4), and Aβ (2^3^=8). Prior to fusion, we upsampled the predictions generated at lower resolutions (GM and LS) to match the native resolution of the vessel and Aβ masks. We combined all predictions using a logical OR operation, producing an overlap map that preserved concurrent classifications at the pixel level. For example, pixels classified as both vessel wall and Aβ deposit were assigned a value of 12. Pixels containing both vessel wall and Aβ labels were subsequently classified as Aβ immunohistochemistry (IHC)-positive vessels. Regions not segmented by either the LS or GM models were implicitly assigned to WM. This rule-based post-processing step enabled explicit separation of vascular Aβ deposits from parenchymal Aβ plaques and from Aβ IHC-negative vessels. The sequential application of models and fusion of their outputs into a composite, color-coded segmentation mask is illustrated in Figure 9.

**Figure 9.**
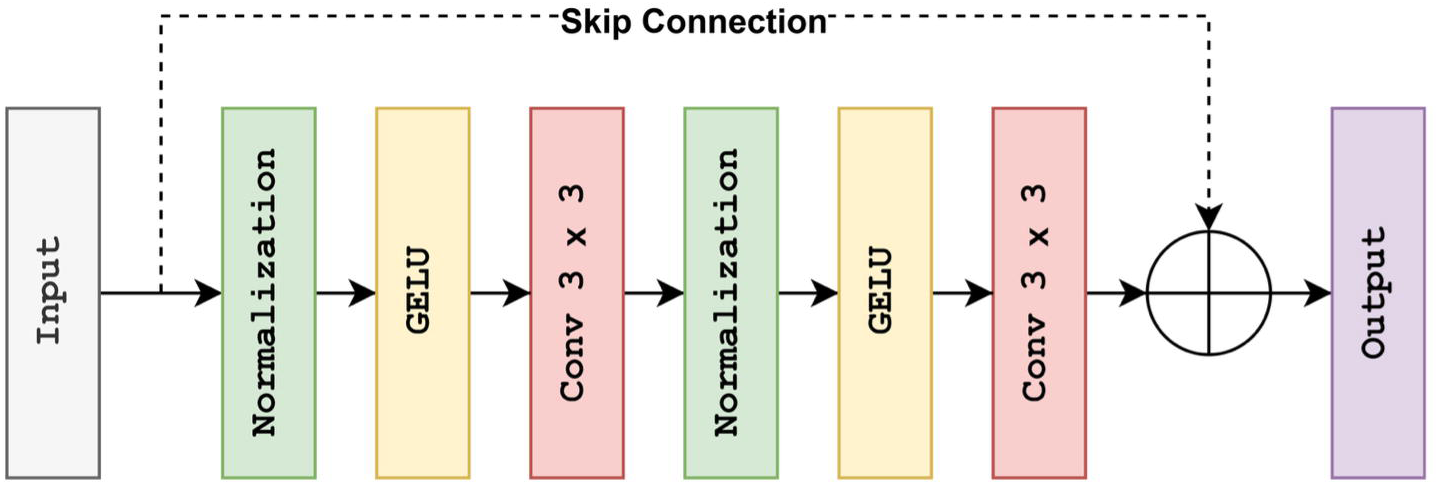
Sequential multi-model inference and composite mask generation. An input image tile is processed independently by four trained binary models targeting specific structures: vessel walls (green), Aβ deposits (red), gray matter (yellow), and leptomeninges (magenta). The resulting individual segmentation maps are subsequently fused to generate a final multi-class composite mask, with overlapping vessel wall and Aβ regions labeled as vascular Aβ (blue). Lepto, leptomeninges.

### 3.10 Vessel instance extraction from composite masks

We derived all vascular morphometrics directly from the composite, color-coded segmentation masks, which encoded vessel walls, Aβ deposits, LS, GM, and WM. All downstream measurements were computed deterministically from these masks and did not require access to the original histology images. We performed connected-component labeling on the vessel wall class and extracted contour hierarchies using OpenCV 4.12 to recover nested anatomical structures. We defined a valid vessel instance as an inner lumen contour fully enclosed by a corresponding outer vessel wall contour. We excluded candidates that failed predefined geometric validity criteria, such as missing enclosure or implausible contour geometry.

### 3.11 Vessel morphometry and wall thickness

We quantified vessel geometry and wall thickness using two complementary computational algorithms, illustrated in Figure 10.

**Figure 10.**
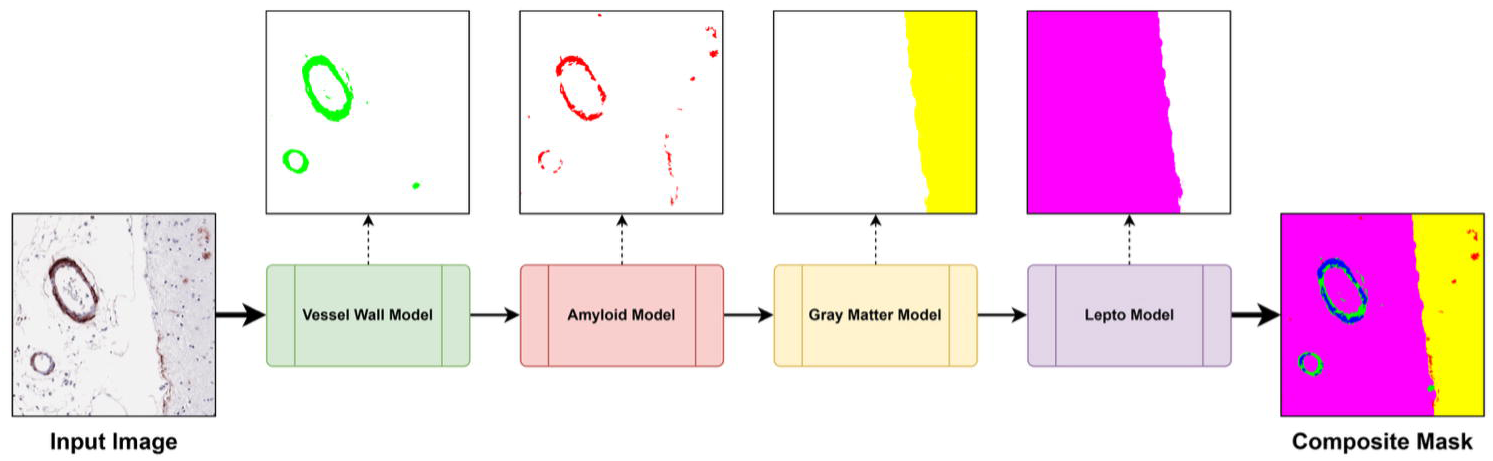
Vessel geometry and wall thickness measurements. A schematic vessel illustrates the lumen centroid *C* (black dot). Radial sampling is shown by the red ray (r_i_) extending from the centroid to the outer vessel wall, defining inner diameter (d_inner_), outer diameter (d_outer_), and radial wall thickness (T_radial_). Tangent-normal measurements are shown at lumen boundary point P_i_, where the local tangent line (t) is shown in green and its perpendicular vector is shown in blue. Wall thickness measured along this normal direction is denoted as T_tangent_ (blue segment).

#### 3.11.1 Radial sampling-based measurements

We first applied a centroid-based radial sampling approach. We calculated the lumen centroid C(x(,y() as the arithmetic mean of the *N* discrete coordinates defining the lumen boundary:

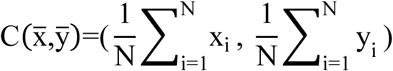

From this centroid, we cast 120 rays at uniform angular intervals (3°) over a full 360° range. Along each ray, we measured the Euclidean distance from *C* to the lumen boundary and to the outer vessel wall boundary (r_i_). We defined the radial wall thickness T_radial_ as the difference between these two scalar distances (Figure 10). To estimate vessel diameters, we summed the radii of antipodal (opposite) rays. We then computed the inner-to-outer diameter ratio (a variation of the classic variable g-ratio) to serve as a dimensionless metric of relative wall thickening.

#### 3.11.2 Tangent-normal sampling-based measurements

In this approach, we measured vessel wall thickness by sampling points along the boundary of the lumen. At each lumen boundary point P_i_, we computed the local tangent vector *t* and computed the outward normal vector as the perpendicular direction to the tangent (Figure 10). We then calculated the normal wall thickness (T_tangent_) by measuring the Euclidean distance along this normal vector from the inner lumen boundary to its intersection with the outer vessel wall. This approach captures local thickness variations independent of the vessel center, providing robust measurements for elongated or deformed lumens.

#### 3.11.3 Vessel roundness

We quantified vessel roundness using a radial uniformity metric derived from the same centroid-based radial sampling framework described above. For each vessel, we summarized the variability of radial distances from the lumen centroid to the outer vessel wall across angular directions. Roundness was defined as:

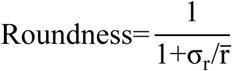

where *r̅* is the mean radial distance and *σ_r_* is the standard deviation of radial distances. This dimensionless metric decreases with increasing radial variability and reflects deviations from circular vessel geometry due to eccentricity or irregular wall morphology.

### 3.12 Quantification of vascular Aβ burden and spatial distribution

We quantified vascular Aβ burden as the proportion of Aβ-labeled pixels within the segmented vessel wall. To characterize the peri-vascular spatial distribution of the deposits, we performed a concentric annular analysis. We generated a single ring-shaped region of interest (ROI) by morphologically dilating the outer vessel wall boundary to a fixed radial distance of 75 µm. This distance was adopted from a prior neuropathological study that defined the perivascular compartment as a fixed-radius region extending from the tunica externa for quantification of Aβ signal (Fastenau et al., 2025).

Within this annulus, we computed the Aβ density, defined as the ratio of Aβ-labeled pixels to the total pixel count of the region.

To evaluate whether vascular Aβ deposition was circumferential or focal, we partitioned the vessel wall into angular sectors aligned with the radial sampling framework. We classified vessels as circumferentially affected when Aβ signal exceeded a 50% occupancy threshold across all sectors. This classification distinguished uniformly encircling deposition from segmental or patchy involvement.

### 3.13 Tissue context assignment

We determined the anatomical localization of each vessel by performing a majority vote of the pixel-level tissue labels within the vessel’s bounding box. We utilized this spatial context to conditionally filter the derived morphometric features. Specifically, we retained perivascular annular measurements for parenchymal vessels (GM) but set these feature values to zero for leptomeningeal vessels.

## 4 Results

### 4.1 Segmentation performance and model comparison

#### 4.1.1 Segmentation performance on the internal development cohort

We evaluated segmentation performance for vessel walls, Aβ deposits, GM, and LS using five-fold cross-validation on the internal development cohort. We compared standard U-Net, DA-ResUNet, and Swin U-Net using Dice coefficient, recall, and precision. Mean values and standard deviations (SD) across folds are reported in Table 7.

**Table 7.**
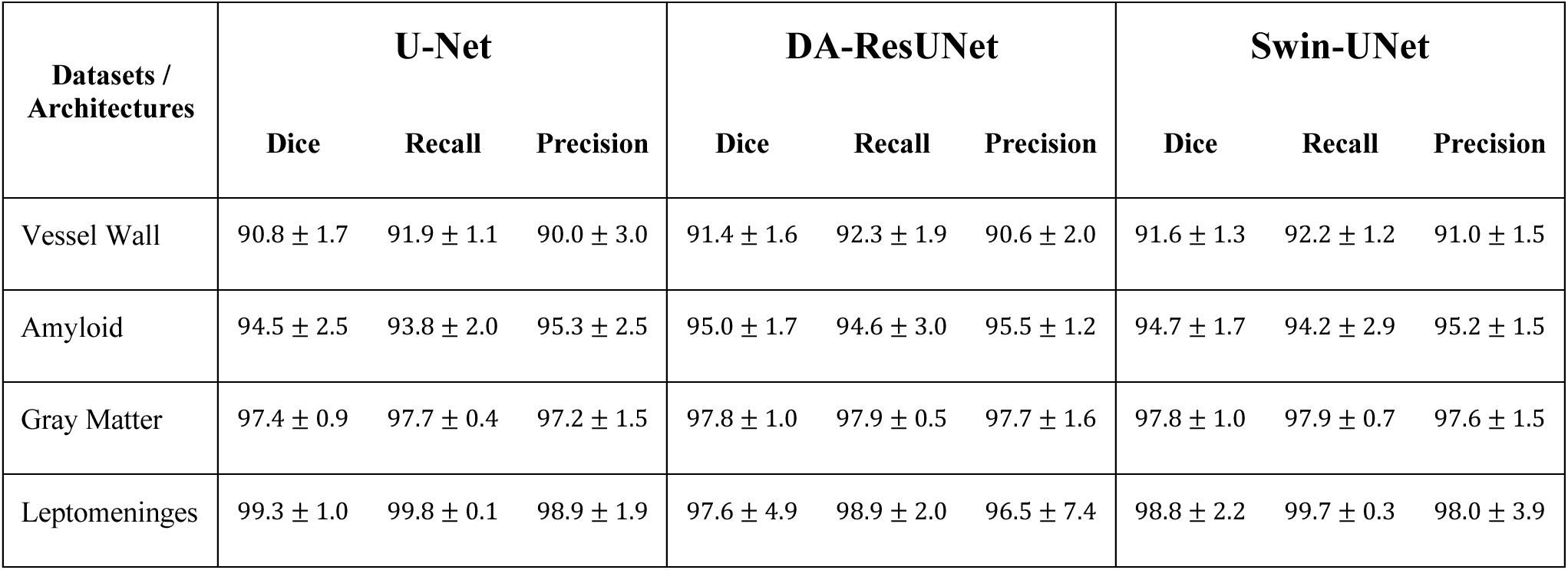
Comparison of segmentation performance metrics (*mean ± SD*) on the internal development cohort.

All three architectures achieved high segmentation fidelity, with Dice scores exceeding 90% for all tissue compartments. The Swin-UNet demonstrated the highest performance for vascular segmentation, achieving a Dice score of 91.6±1.3%, compared to 91.4±1.4% for the DA-ResUNet and 90.6±1.7% for the Standard U-Net.

For fine-grained pathological features, the DA-ResUNet exhibited a slight advantage. In the amyloid segmentation task, the DA-ResUNet achieved the highest Dice score (95.0±1.7%) and Precision (95.5±1.2%). Swin-UNet showed comparable but slightly lower performance with a Dice score of 94.7±1.7% and Precision of 95.2±1.2%.

Notably, the Standard U-Net remained highly competitive, particularly for large-scale structural regions. For LS, it achieved the highest Dice score of (99.3±1.0%), outperforming both the Swin-UNet (98.8±2.2%) and the DA-ResUNet (97.6±4.9%). This indicates that for distinct, macroscopic tissue boundaries, the added complexity of attention mechanisms may not be necessary for optimal segmentation. For GM, both advanced architectures performed identically (97.8 ± 1.0%) slightly surpassing the Standard U-Net (97.4±0.9%).

Visual inspection of the segmentation masks confirmed that both the DA-ResUNet and Swin-UNet produced sharper boundaries than the Standard U-Net (Figure 11). We also note that the reference annotations are not error free, and in some cases the model predictions corrected minor inconsistencies or imprecision in the ground truth delineations.

**Figure 11.**
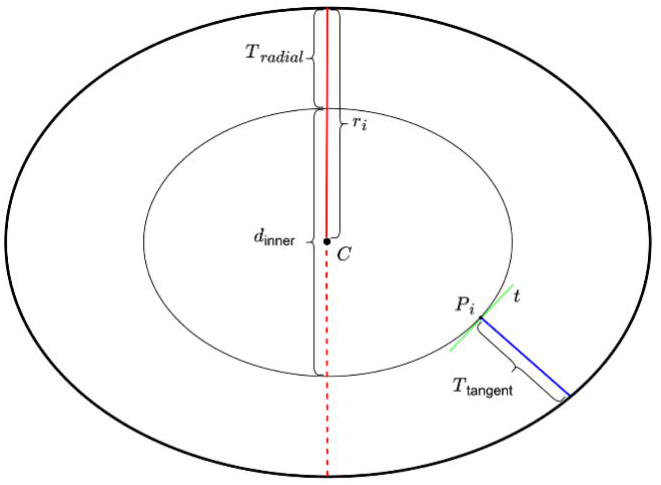
Example segmentation outputs across models. Representative examples showing input images, ground truth annotations, and corresponding segmentation outputs from the standard U-Net, DA-ResUNet, and Swin-UNet for vessel walls, Aβ deposits, leptomeninges, and gray matter. In these examples, DA-ResUNet and SwinUNet show improved delineation, particularly for Aβ deposits and gray matter.

Based on this performance and its computational efficiency, we selected DA-ResUNet as the segmentation backbone for all subsequent vessel-level and morphometric analyses.

#### 4.1.2 Generalization to an independent external cohort

We evaluated model generalization using tiles extracted from the UKY cohort, which were not used during training or validation. For each cohort, we tested around 500 tiles per target class sampled from 10 WSIs. Table 8 summarizes the performance of the three architectures on this unseen data.

**Table 8.**
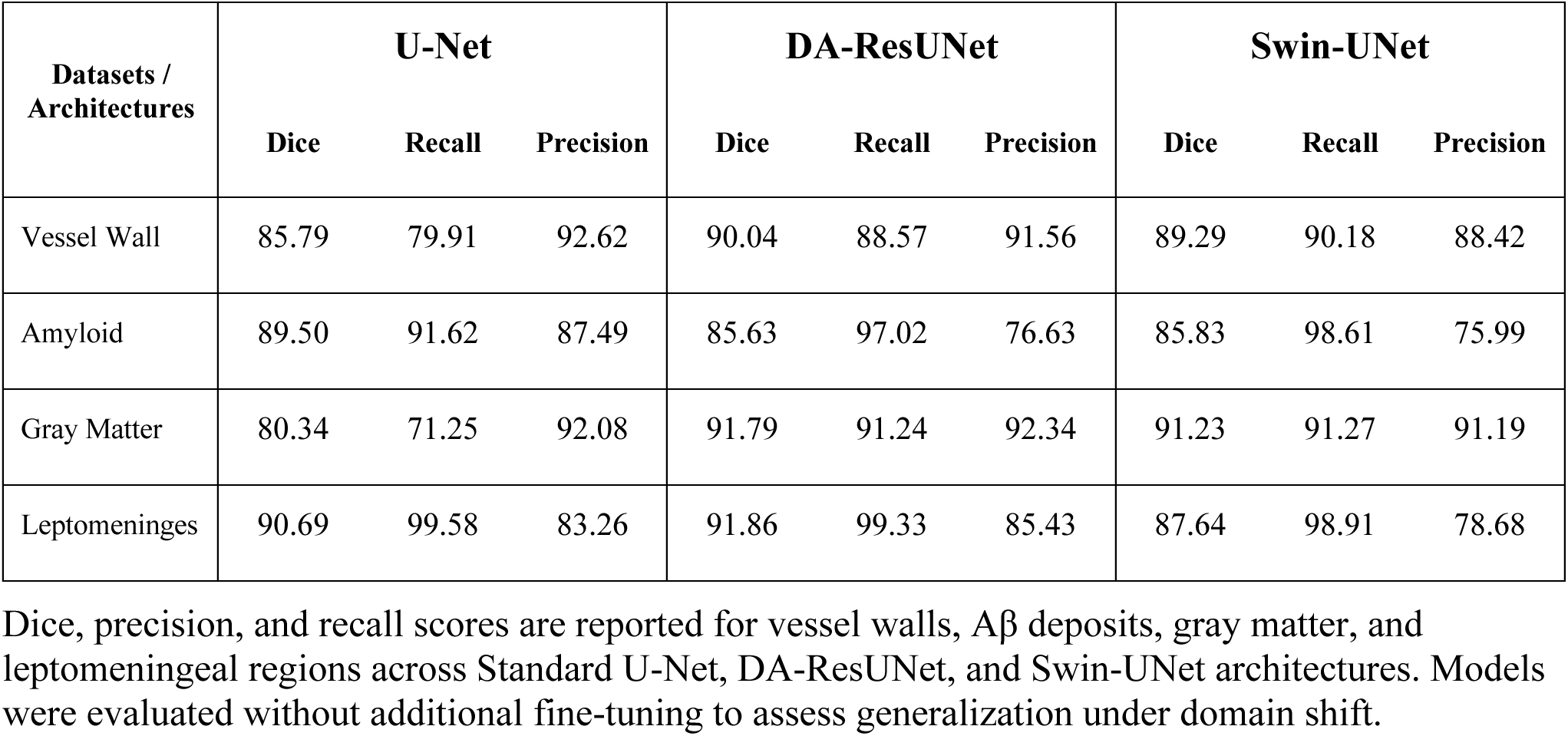
Segmentation performance on independent external cohorts.

For vessel wall segmentation, the DA-ResUNet achieved the highest Dice score (90.04%) and preserved sensitivity more effectively than the Standard U-Net, whose performance declined primarily due to reduced recall (79.91%). This performance gap was more pronounced in GM segmentation, where the Standard U-Net showed limited generalization (Dice: 80.34%), while both the DA-ResUNet (91.79%) and Swin-UNet (91.23%) maintained high accuracy. In the Aβ segmentation task, the Standard U-Net achieved the highest Dice score (89.50%). In contrast, the attention-based architectures exhibited higher recall (>97%) but lower precision, suggesting increased sensitivity to ambiguous signal and a tendency toward over-segmentation in unseen data. Finally, for leptomeningeal segmentation, the DA-ResUNet outperformed both alternatives (Dice: 91.86%), indicating improved architectural stability for large, well-defined tissue compartments under domain shift.

### 4.2 Detection and quality filtering of individual blood vessels

The pipeline successfully detected individual blood vessels across GM, WM, and LS based on the presence of a continuous vessel wall enclosing a clearly defined lumen. Detected vessels exhibited anatomically plausible geometry and were suitable for downstream morphometric analysis.

Structures that did not meet these criteria were automatically excluded, most commonly due to fragmented or discontinuous vessel walls that prevented reliable identification of an enclosed lumen. Furthermore, we applied a minimum lumen area threshold of 20µm^2^ to exclude candidate vessels with poorly resolved lumens at the available image resolution, while retaining valid capillary-scale vessels. This defined minimum threshold is modifiable by users. Representative examples of successfully detected vessels and excluded non-vascular artifacts are shown in Figure 12.

**Figure 12.**
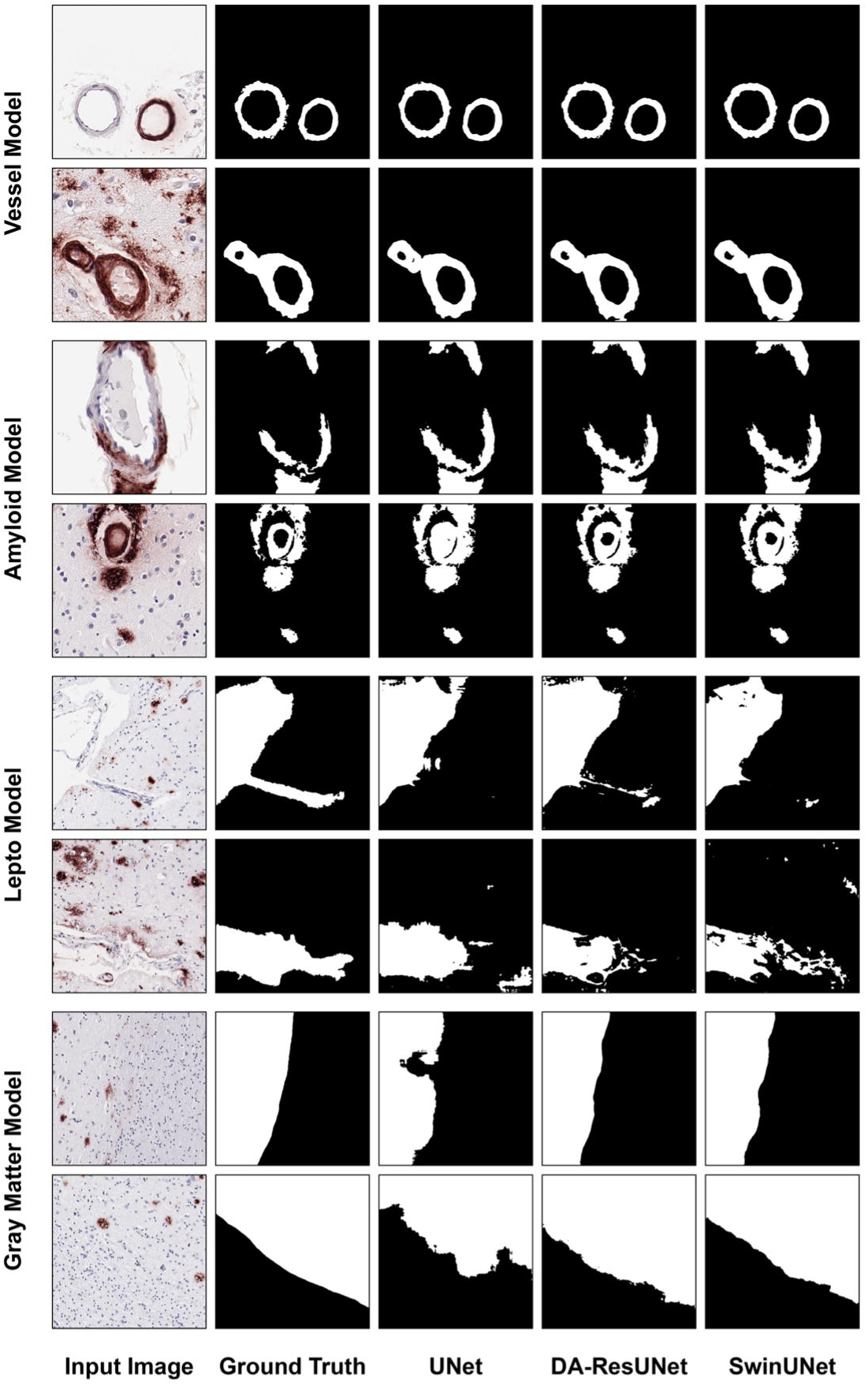
Representative examples of vessel detection and exclusion criteria. (A) Vessel segmentation in the leptomeninges, indicated by the magenta background in the composite masks. (B) Vessel segmentation in the cortex, indicated by the yellow background. The panels display the original input images, the generated composite masks, and the final detected vessels. Bounding boxes enclose complete vessels included in the analysis, while arrowheads point to incomplete vessels, artifacts or tangentially cut vascular structures without a lumen that were excluded. Scale bars = 50μm.

### 4.3 Validation of geometric characterization and CAA identification

Vessel-level identification of CAA was performed by spatial co-localization of vessel wall and amyloid segmentation masks. For each detected vessel, the pipeline computed inner and outer diameters, the inner-to-outer diameter ratio, vessel roundness, and wall thickness using both radial (T_radial_) and tangent-normal (T_tangent_) measurements. Aβ burden was quantified within the vessel wall and within a perivascular ring-shaped annulus to capture Aβ extending beyond the vessel boundary. Based on the angular distribution of Aβ signal, vessels were further classified as circumferentially affected or focally involved. Representative examples of these vessel-level measurements are shown in Figure 13. As a preliminary validation of the pipeline, occipital cortex sections showed higher vessel amyloid burden, more circumferential vessel involvement, and more Aβ-positive capillary vessels than frontal cortex sections.

**Figure 13.**
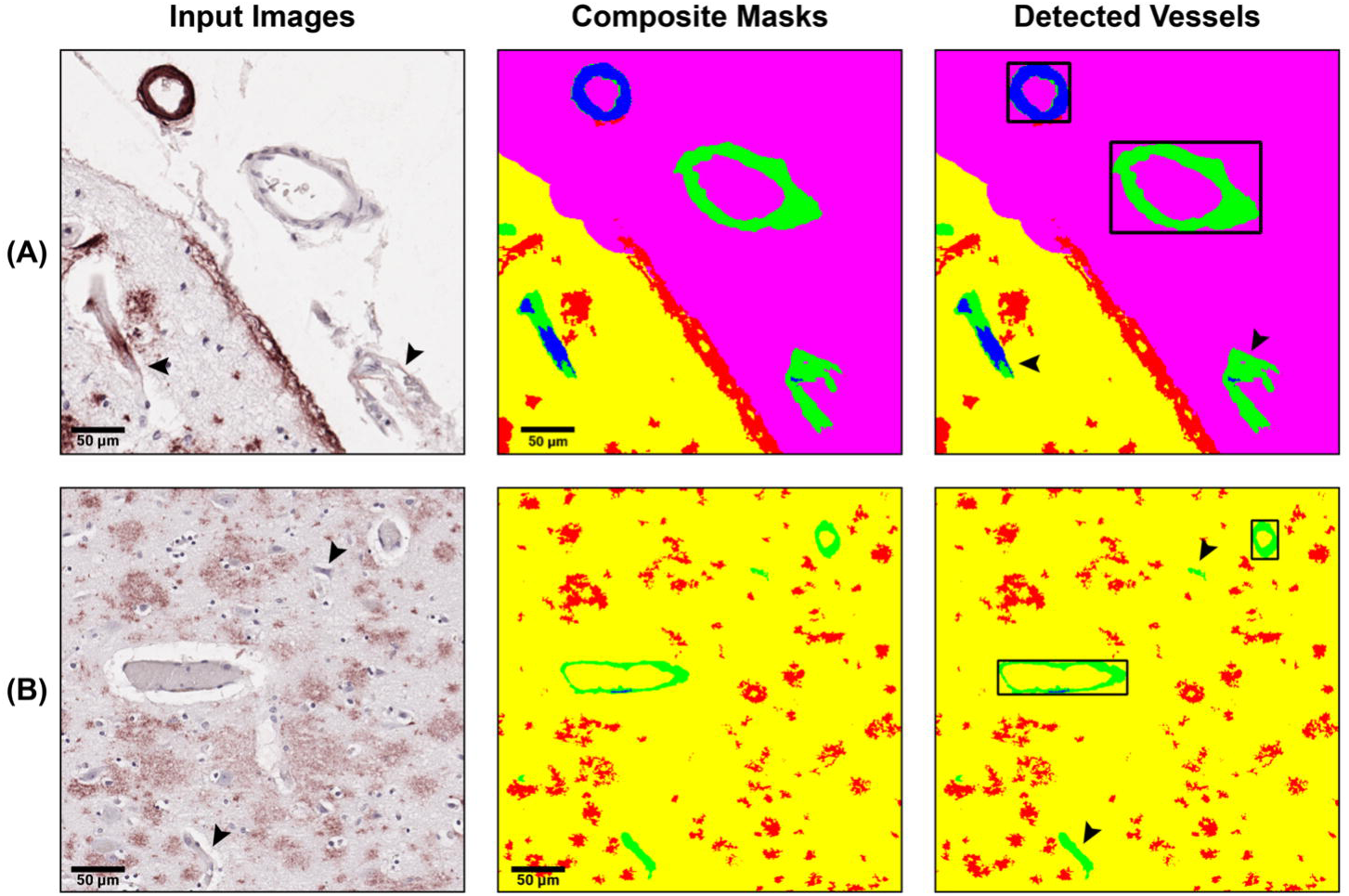
Example outputs generated by the automated vessel-level analysis pipeline. (A) Panel A shows a cortical parenchymal vessel with dyshoric changes, and (B) panel shows a leptomeningeal vessel. All measurements displayed were produced automatically by the pipeline. For each vessel, the pipeline reports the minimum, maximum, and mean values of inner diameter, outer diameter, and wall thickness. Vessel roundness is quantified from the variability of radial distances measured from the lumen centroid to the outer vessel wall, with lower radial dispersion indicating a more circular geometry. Vascular Aβ burden is quantified as the percentage of Aβ IHC-positive pixels within the vessel wall. Circumferential involvement is determined based on angular coverage of Aβ signal around the vessel wall. Perivascular Aβ deposition is quantified as percent Aβ burden within a fixed ring-shaped annulus surrounding the outer vessel wall. Scale bars = 50μm. * Aβ in ring is not defined for leptomeningeal vessels.

## 5 Discussion

In this study, we developed and evaluated a DL-based framework for automated, vessel-level analysis of cerebrovascular pathology in WSIs of human post-mortem brain tissue. The pipeline combines DL-based segmentation with deterministic geometric feature extraction to enable quantitative assessment of blood vessel morphology and vascular Aβ involvement. We compared convolutional and transformer-based segmentation architectures, assessed generalization across independent external cohorts, and demonstrated reliable detection of individual vessels across GM, WM, and LS. By deriving all measurements directly from composite segmentation masks, the approach supports scalable and reproducible vessel-level quantification without reliance on semi-quantitative grading systems. This framework provides a generalizable foundation for computational analysis of CAA and related vascular pathologies in WSIs.

We evaluated three segmentation architectures, standard U-Net, DA-ResUNet, and Swin-UNet, to examine trade-offs between segmentation accuracy, generalization, and computational efficiency in WSI analysis. The Swin-UNet showed nominally higher performance for vascular segmentation, which is consistent with the ability of transformer-based self-attention to model long-range spatial dependencies and preserve continuity in elongated structures beyond the local receptive fields of conventional convolutional networks (Thisanke et al., 2023). Such global context is advantageous for maintaining vessel integrity across large fields of view. However, this representational capacity comes at a substantial computational cost. Self-attention mechanisms remain more resource-intensive than convolutional operations, even when constrained through window-based and shifted-window attention as implemented in Swin Transformers (Liu et al., 2021; Palanisamy et al., 2025). In our implementation, this resulted in a markedly higher computational overhead: while the DA-ResUNet processed a typical WSI (approx. 80,000 x 80,000 pixels) in approximately 4-5 minutes, the Swin-UNet required nearly 8-10 minutes under identical hardware conditions. This 2-fold increase in processing time implies that despite comparable theoretical FLOPs, the memory access patterns of the Transformer architecture create a significant bottleneck, potentially limiting its viability for high-throughput clinical workflows where rapid case turnover is essential.

In contrast, the DA-ResUNet achieved a more favorable balance between performance and efficiency. Its residual design improves gradient flow and feature reuse, while the integration of channel and spatial attention selectively emphasizes informative regions and suppresses background noise. This architectural bias is well suited for detecting sparse, fine-grained pathological features such as vascular Aβ deposits, where subtle intensity and texture cues are critical. Similar observations have been reported in other medical imaging domains, where attention-augmented residual U-Net variants improved lesion segmentation by expanding the effective receptive field and enhancing multi-scale feature integration without incurring the computational burden of transformer backbones (Sen et al., 2025; Shah & Kang, 2023; Zhang et al., 2025). For example, attention-enhanced residual U-Net models have demonstrated improved performance over standard U-Net architectures in breast ultrasound lesion segmentation, particularly for small or poorly defined targets, while maintaining computational efficiency suitable for clinical workflows (Zhang, 2025).

Evaluation on an independent external cohort revealed differences in model robustness under domain shift that reflect architectural inductive biases. The Standard U-Net showed reduced stability in heterogeneous tissue regions, likely due to reliance on low-level appearance features such as staining intensity and local texture, which vary across scanners and sites. In contrast, the DA-ResUNet and Swin-UNet maintained more consistent behavior, suggesting that attention mechanisms promote greater reliance on structural and spatial features rather than pixel-level appearance. Inter-site variability is a critical issue in multi-center histopathology because differences in tissue processing, staining, and digitization can introduce large appearance shifts that are unrelated to biology and often degrade model generalization (Asadi-Aghbolaghi et al., 2024; Tellez et al., 2019).

Unsurprisingly, the models showed different training behaviors. The Swin-UNet converged faster during early training but began to overfit at later epochs, as validation loss increased while training loss continued to decrease. This behavior is consistent with previous studies showing that transformer-based models, due to their high model capacity, can overfit when training data are limited (Touvron et al., 2021). In contrast, the DA-ResUNet showed more stable training over longer schedules, while the standard U-Net required more epochs to converge.

We next evaluated the sequential use of multiple binary segmentation models and the generation of a composite, color-coded segmentation mask. This design avoided the class imbalance challenges that commonly affect multi-class segmentation models in histopathology (Sudre et al., 2017). Training one model per target minimized class competition and allowed explicit control of model-specific errors. We adjusted the loss function asymmetrically, penalizing FPs more strongly for the Aβ model to limit spurious CAA detection, and penalizing FNs more strongly for the vessel wall model to favor complete vessel identification. The primary limitation of this approach is increased inference time compared with a single multi-class model.

The pipeline’s vessel detection prioritizes geometric validity over maximal sensitivity. By requiring a continuous vessel wall and a detectable lumen, the framework effectively excluded nonvascular artifacts and isolated false-positive pixels, which is critical for reliable downstream morphometric analysis. Although vessels with fragmented walls or small lumens may be missed, this trade-off is acceptable in a vessel-centric analysis where false positives can propagate substantial bias into geometric measurements. The deterministic quality filtering, together with asymmetric loss weighting during training, enabled consistent vessel identification while maintaining biological plausibility.

Vessel morphometry presents additional challenges due to sectioning artifacts and shape variability. Tangentially cut vessels or vessels with highly asymmetric geometry can distort diameter and thickness estimates when measured using purely radial assumptions. In the analyses of these structures, large discrepancies between opposing radial measurements serve as indicators of geometric asymmetry. The pipeline addresses this variability through rule-based handling, including conservative diameter estimation and the use of tangent-normal thickness measurements when radial estimates become unreliable. Measuring wall thickness perpendicular to the lumen boundary reduces sensitivity to lumen eccentricity and better captures focal wall remodeling. These complementary strategies allow robust geometric quantification across diverse vessel shapes while preserving interpretability and minimizing bias introduced by histological sectioning artifacts.

Quantification of vascular Aβ burden captured both intramural deposition and perivascular extension. Estimating Aβ fraction within the vessel wall provided a direct measure of vascular involvement, while analysis of a ring-shaped perivascular region enabled detection of Aβ extending into the adjacent neuropil. This pattern is characteristic of dyshoric CAA, where Aβ deposition spreads beyond the vessel wall into surrounding tissue (Richard et al., 2010). Incorporating this spatial context allows the pipeline to distinguish Aβ immunoreactivity that is confined to the vessel wall from parenchymal Aβ deposition that extends into the surrounding tissue. Unlike conventional semiquantitative grading, which is based on visual assessment of a limited number of vessels, this framework provides exact vessel-level measurements of vascular Aβ burden, perivascular Aβ burden, and vessel geometry across all analyzable vessels in a section. For this reason, the pipeline is best viewed as a quantitative complement to routine neuropathologic assessment. In particular, it can serve as a research tool for large-scale, reproducible extraction of vascular features that are not practical to obtain manually. These measurements can support identification of vessel-level patterns, clustering of CAA phenotypes, and comparisons across brain regions, cohorts, and disease states. For example, in the present cohort, the pipeline identified more circumferentially affected vessels, more Aβ-positive capillary vessels, and higher average vascular amyloid burden in the occipital cortex than in the frontal cortex. These observations are consistent with prior biologic studies showing that CAA is more severe in the occipital cortex (Attems et al., 2005).

The current study has several limitations. Vessel detection relied on strict geometric rules that favor specificity over sensitivity. At vessel bifurcation sites, where two lumens may be present, or in regions where adjacent vessel walls are in close contact, the pipeline retained only the vessel with the larger lumen. Similarly, even small discontinuities in the vessel wall could lead to exclusion of otherwise valid vessels. While this strategy effectively removed nonvascular artifacts, it may result in missed detections in complex vascular regions. In addition, although the models demonstrated good generalizability across the institutions we tested, larger variations in staining or scanner characteristics could hinder performance. Of note, our models were developed for Aβ immunostained sections only. Other staining protocols, such as hematoxylin and eosin, require separate evaluation and potential fine-tuning. We also did not perform formal cross-magnification validation. Vessel wall and Aβ models were trained and validated at 40X, whereas GM and LS models were trained and validated at 10X. Lower effective resolution may reduce sensitivity for very small vessels, particularly capillaries. The sample size was also limited, and broader patient-level and institution-level cohort diversity may reveal additional variability in vessel morphology and Aβ deposition patterns. Quantification of perivascular Aβ was based on a fixed ring-shaped annulus at a predefined distance from the vessel wall, which may not capture parenchymal Aβ in cases with enlarged perivascular spaces. Finally, given that a single Aβ IHC-positive pixel within a vessel wall is sufficient to classify a vessel as CAA-positive in our implementation, this framework favors sensitivity and may overestimate vascular involvement in borderline cases.

Future work will address these limitations by expanding training data to include larger cohorts and greater staining variability, which may improve robustness and refine vessel- and Aβ-level characterization. We also plan to provide model checkpoints trained at different resolutions so that users can select the most appropriate model for their scanning workflow. Additional rules or probabilistic criteria could be incorporated to better handle bifurcations, fragmented walls, and ambiguous vessel geometries. The modular design of the pipeline also allows integration of additional models to detect related pathologies, such as microhemorrhages, infarctions, and immune cell infiltration, enabling more comprehensive characterization of CAA-associated tissue changes. Ultimately, the resulting quantitative vessel-level features could support definition of CAA endophenotypes and facilitate analysis of their relationships with clinical presentation, disease progression, and genetic factors. Beyond CAA, the same strategy may also support quantitative study of other small-vessel diseases, including arteriolosclerosis.

In conclusion, we present a modular DL framework for vessel-level analysis of CAA in WSIs. By combining target-specific segmentation with deterministic geometric measurements, the pipeline enables reproducible quantification of vascular structure and Aβ immunoreactivity involvement at scale. The design emphasizes interpretability, robustness, and flexibility, providing a foundation for future extensions to additional vascular pathologies and integrative analyses linking imaging-derived features with clinical and genetic data.

## 6 Abbreviations

Aβ: amyloid-beta
AD,sa: Alzheimer’s disease
ADRC: Alzheimer’s Disease Research Center
AHA: American Heart Association
BCE: binary cross-entropy
CAA: cerebral amyloid angiopathy
CBAM: Convolutional Block Attention Module
CERAD: Consortium to Establish a Registry for Alzheimer’s Disease
CNN: convolutional neural network
CORID: Committee for Oversight of Research and Clinical Training Involving Decedents
CUDA: Compute Unified Device Architecture
DA-ResUNet: dual-attention residual U-Net
DL: deep learning
FLOPs: floating point operations
FN: false negatives
FP: false positives
GELU: Gaussian Error Linear Unit
GFLOPs: giga floating point operations
GM: gray matter
GPU: graphics processing unit.

## 8 Author Contributions

**HT:** Conceptualization, Methodology, Validation, Investigation, Writing - original draft. **AB:** Methodology, Writing - review & editing. **DRJ:** Methodology, Writing - review & editing. **JAC:** Methodology, Writing - review & editing. **MN:** Data curation, Resources. **VKCB:** Resources. **PTN**: Resources, Writing - review & editing. **TMP:** Conceptualization, Methodology, Supervision, Data curation, Resources, Writing - review & editing. **JK:** Conceptualization, Methodology, Supervision, Validation, Resources, Writing - review & editing.

## Acknowledgments

This research was supported in part by the University of Pittsburgh Center for Research Computing through the resources provided. Specifically, this work used the HTC cluster, which is supported by NIH award number S10OD028483.

